# Above- and below-ground biodiversity responses to the prolonged flood pulse in central-western Amazonia, Brazil

**DOI:** 10.1101/2021.04.28.441795

**Authors:** Yennie K. Bredin, Laura L. Hess, Andressa B. Scabin, Micah Dunthorn, Torbjørn Haugaasen, Carlos A. Peres, Henrik R. Nilsson, Alexandre Antonelli, Camila D. Ritter

## Abstract

Amazonia encompasses forests that grow in areas that are periodically inundated by overflowing rivers. The inundation depth and duration vary according to the slope of the terrain, creating a flooding gradient. This gradient directly affects the biota, but the effect on soil organisms remains elusive. Here, we use DNA metabarcoding to estimate prokaryote and eukaryote diversity from soil and litter samples in a seasonally flooded forest and its adjacent unflooded forest in central-western Amazonia using 16S and 18S gene sequences, respectively. We characterize the below-ground diversity and community composition based on Amplicon Sequence Variants (ASVs) along the flooding gradient. We test for the relationship of soil biota with the flooding gradient, soil properties and above-ground woody plant diversity. The flooding gradient did not explain below-ground biodiversity. Nor was the below-ground diversity explained by the above-ground woody plant diversity. However, we found taxonomic groups not previously reported in Amazonian seasonally flooded forests. Also, the flooding gradient and woody plant diversity did, in part, explain the community composition of soil bacteria. Although the effects of the flooding gradient, soil properties and above-ground woody plant diversity is hard to quantify, our results thus indicate that flood stress could influence below-ground bacterial community composition.

## 1. Introduction

Amazonia comprises the largest continuous tropical rainforest in the world. Accounting for only 3.6% of the terrestrial global surface, Amazonia harbours 10% of the world’s known biodiversity (Maretti, 2014) and potentially hosts the largest Linnaean biodiversity knowledge deficit on Earth (Moura and Jetz, 2021). Amazonia is heterogeneous and encompasses several distinct environments. These include tropical rainforests known as terra firme, non-forested areas, such as the edaphic open areas associated with white sand soils, and seasonally flooded forests (Myster, 2016). Seasonally flooded forests grow in areas that are periodically inundated by overflowing rivers, lakes and perennial streams (Prance, 1996). These forests are characterized by low taxonomic diversity compared to terra firme forests (Haugaasen and Peres, 2006; Myster, 2016; ter Steege and Hammond, 2001). However, they have a characteristic fauna and flora often restricted to these environments (Myster, 2016; Ramalho et al., 2016). At least 9% of the Amazon basin is formed by seasonally or permanently flooded forests (Hess et al., 2015), which are crucial for the maintenance of biodiversity and climatic dynamics in the region (Castello and Macedo, 2016).

Two determinants are decisive for the extent of periodically flooded forests in Amazonia. The first is the uneven annual distribution of rainfall. In most parts of Amazonia, the rainy season is followed by a drier period lasting several months, but this is not synchronous across the basin. The second is the topography of the Amazon basin and its low-lying floodplains. Combined, these factors lead to an annual rise in fluvial discharge which causes an enormous flood pulse (Junk, 1989; Kubitzki, 1990) and gives rise to an aquatic and a terrestrial phase in the flooded areas. The inundation depth and duration of the flood waters vary according to the slope of the terrain and the volume of the rivers that flood the landscape (Assis et al., 2015; Wittmann et al., 2010). This creates a gradient in flood depth and duration from low-lying areas flood to greater depths for longer periods of time to areas higher up in the terrain that flood for shorter periods. This gradient directly affects the biota, generating thresholds for species establishment (Petit and Hampe, 2006). Additionally, the physical and chemical properties of the waters also affect the distribution of biota in inundated areas (Prance, 1979).

In the Amazon basin, seasonally flooded forests can be classified into two major types according to the hydro-chemical characteristics of the rivers that flood them (Assis et al., 2015; Haugaasen and Peres, 2006; Myster, 2016; Prance, 1979). Whereas eutrophic várzea forests are flooded by nutrient-rich white-water rivers originating in the Andes, oligotrophic igapó forests are inundated by nutrient poor, black- or clear-water rivers (Ríos-Villamizar et al., 2020). Thus, fluvial geochemistry determines the physical properties of substrate, such as moisture retention and hydraulic conductivity, accumulation of organic matter, nutrient availability and soil biota (Parolin et al., 2004). It has been demonstrated that changes in above-ground species richness and composition in seasonally flooded forests can occur due to the physicochemical characteristics of the water (Myster, 2016) and/or flood depth (Julião et al., 2018). Few studies have evaluated this difference in soil biota (Ritter et al., 2019b), and to our knowledge no study has yet examined the influence of the flooding gradient on seasonally flooded forest soil biodiversity.

Soil biota represent a large reservoir of terrestrial biodiversity and provide fundamental ecosystem services that are key to the functionality of terrestrial ecosystems (Bardgett and Van Der Putten, 2014; Pereira et al., 2018; Pietramellara et al., 2002). For instance, larger soil invertebrates are responsible for processing large amounts of detritus and make it available to other organisms (García□Palacios et al., 2013; Hättenschwiler and Gasser, 2005). Similarly, micro-organisms are essential for nutrient cycling (Delgado-Baquerizo et al., 2020), and ectomycorrhizal fungi underlie ecosystem processes such as soil carbon cycling (Johnson et al., 2016). Yet, soil biodiversity remains elusive and has been neglected in many global biodiversity assessments and policies (Cameron et al., 2019; Ritter et al., 2017). This omission is undoubtedly related to the scarcity of comprehensive information on soil biodiversity, especially in megadiverse and remote tropical environments, such as Amazonia. Fortunately, molecular approaches, including high throughput sequencing (HTS), such as metabarcoding (Creer et al., 2016), are now able to address many previous obstacles to understanding the diversity and composition of soil communities (Cameron et al., 2019; Ritter et al., 2019b; Tedersoo et al., 2014).

In this study we use a metabarcoding approach to characterize the soil biodiversity along the flooding gradient of an Amazonian várzea landscape. More specifically, we investigate the diversity and composition of soil communities across three flood-levels and explore if, and how, soil biota changes along the flooding gradient. In addition, by comparing the soil communities to the above-ground woody plant community, we examine the degree to which the above- and below-ground biodiversity are congruent. The results are discussed in relation to other studies and interpreted in light of differences experienced by seasonal flooding, soil characteristics and above-ground woody plant diversity. Finally, we draw some general implications to the conservation of Amazonian biota.

## 2. Materials and Methods

### 2.1. Study area

We conducted the study in the Uacari Sustainable Development Reserve (RDS Uacari) and nearby forests along the central reaches of the Juruá River, western Brazilian Amazonia (Fig. 1). The climate of the region is hot and humid with a mean annual temperature of ∼27°C, average annual rainfall of ∼3,679 mm, and a well-defined rainy season from December until May (Hawes and Peres, 2016). We sampled above-ground woody plant communities and below-ground microbial communities at three different flood levels in várzea (VZ) and adjacent upland forest (i.e. terra firme, TF) that does not flood on a seasonal basis. This “unflooded” forest is growing on Pleistocene floodplain sediments (i.e., paleo-várzea sediments; Assis et al., 2015) abandoned by the meandering Juruá River and at higher elevations than the river’ maximum flood level. The várzea communities were sampled during the 2016 and 2017 dry seasons and the terra firme communities were sampled in the 2017 wet and dry seasons.

**Fig. 1.**
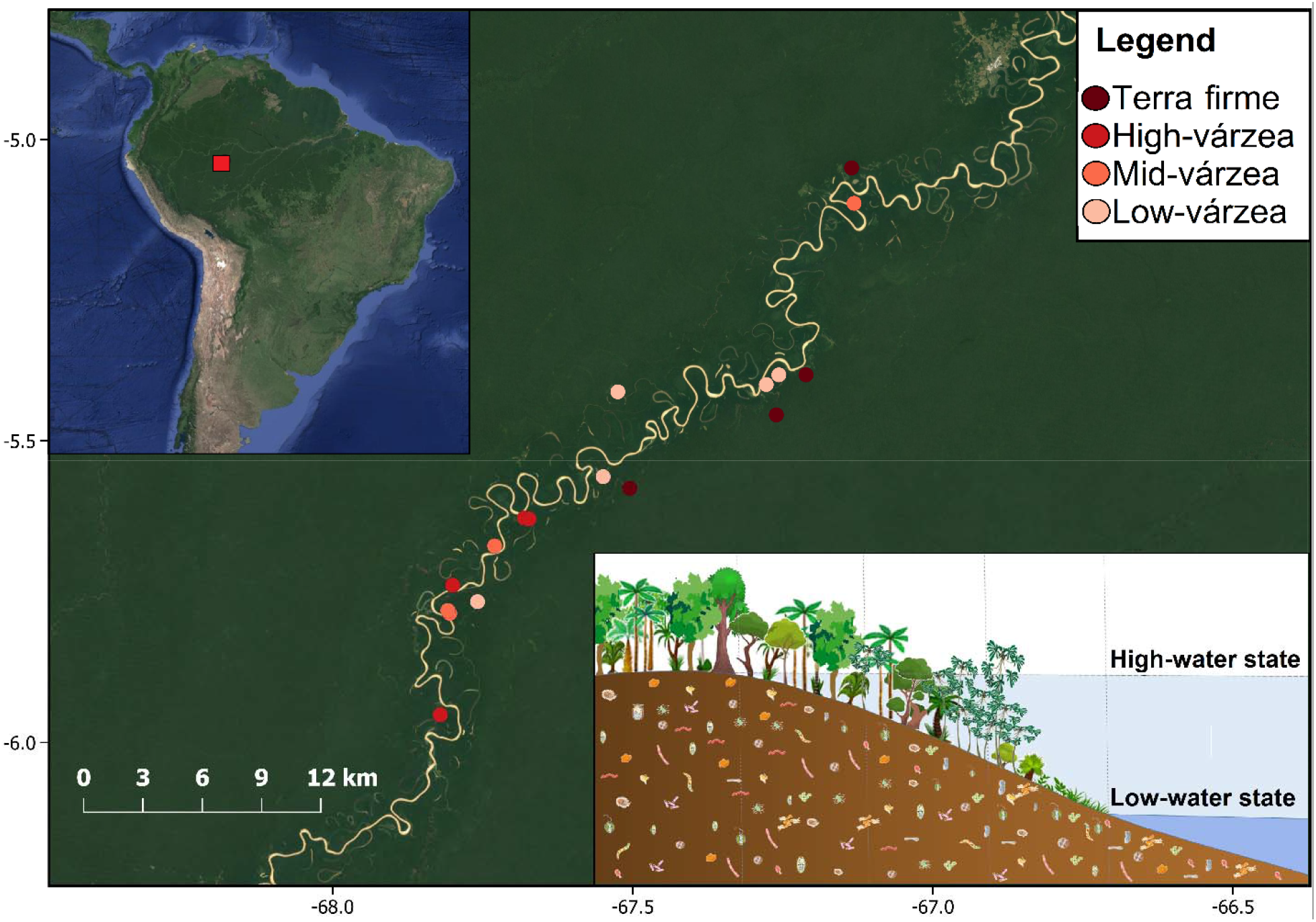
Sampling localities. along the central Juruá River (main map) in the central-western Brazilian Amazon (upper left inset). The lower right inset shows a schematic cross-section of flood levels in the várzea forest, with low- and high-water states separated by the dotted vertical lines. Low-várzea is low-lying and subject to the longest flooding periods (5-12 mo/yr); mid-várzea is subject to intermediate periods of flooding (2-4 mo/yr); and high-várzea is located higher up in the terrain and subject to the shortest flooding periods (0-1 mo/yr). Terra firme forests are beyond the maximum flood levels of rivers and perennial streams. Map created using QGIS3 software (Q. D. Team, 2015).

### 2.2. Determination of the hydro-topographic gradient

To position the plots along the hydro-topographic gradient, we used inundation period mapped with multi-date ALOS-1 PALSAR satellite imagery (Fine-beam mode, resampled to 30 m) freely available from the Alaska Satellite Facility Distributed Active Archive Center (asf.alaska.edu). Water levels at the Porto Gavião gauge on the Juruá River (66.9 W, 4.88 S) were retrieved from Brazil’s Agência Nacional de Águas (ANA; http://www.snirh.gov.br/hidroweb/serieshistoricas) for each of the 28 PALSAR imaging dates between 2007 and 2011 (9-10 dates for each of 3 PALSAR swaths covering the forest plots). The average number of months inundated per year were calculated over the 47-year Gavião river level record (1972-2018). Due to small-scale variability in flood duration even at the 0.1 ha scale, we defined the flooding gradient by approximating the average number of months each plot was flooded annually. Thus, plots were grouped into the following four flood levels: (1) terra firme = not seasonally flooded (n = 6); (2) high-várzea = 0-1 mo/yr, maximum high-water levels < 1.5 m (n = 6); (3) mid-várzea = 2-4 mo/yr, maximum high-water levels = 1-2 m (n = 6); and (4) low-várzea = 5-12 mo/yr, maximum high-water levels ≥ 2 m (n = 4). Flood depth within each plot was determined by measuring the height of visible watermarks left on tree trunks within each plot after the most recent inundation peak. These measurements were made with a measuring tape to the nearest mm.

### 2.3. Above-ground woody plant diversity

We used 0.1 ha floristic plots (100 m x 10 m) placed perpendicular to the main river channel to minimize variability in flood depth and duration within plots. We inventoried woody plant diversity as described in Bredin et al. (2020). Briefly, within each floristic plot, all trees, hemiepiphytes, and palms ≥10cm diameter at breast height (dbh) – as well as all high-climbing woody lianas ≥ 5 cm dbh – were measured and identified. Individuals that could not be determined to species level were sorted to morpho-species or, where applicable, higher taxonomic levels. For the following analyses we only retained floristic data from plots where we also obtained information about substrate biota (n = 18).

### 2.4. Below-ground microbial diversity

To allow for comparisons with other studies of below-ground biodiversity, we used the sampling strategy described in Tedersoo et al. (2014) and Ritter et al. (2019b). Briefly, we superimposed 22 circular plots with a 28 m radius over the floristic plots by matching exactly the midpoints of the circular substrate plot with those of the rectangular floristic plots. Within each circular plot, we randomly selected 20 trees and collected litter and soil samples at the opposite sides of each stem. We first took a litter sample at every sampling point. After removing the leaf litter, we used a soil auger (2.5 cm in diameter) to collect the top 5 cm of the soil. In total, we collected litter and soil at 40 points per plot. The samples were then mixed to provide one composite litter sample and one composite soil sample per plot. For each plot, soil samples were divided into two parts. The first part was sun-dried and transported to the EMBRAPA laboratory in Manaus (Brazil) where physicochemical analyses were performed following standardized procedures (Donagema et al., 2011; Ritter et al., 2018). The second part of the soil samples, as well as the litter samples, were dried with sterilized white silica gel 1–4 mm and transported to the University of Gothenburg, Sweden, for DNA extraction.

### 2.5. DNA extraction and sequencing

For total DNA extraction, we used the PowerMax® Soil DNA Isolation Kit (MO BIO Laboratories, USA) according to the manufacturer’s instructions. We used 10 g (dry weight) from all soil samples and 15 ml of the litter samples (corresponding to 3–10 g of dry weight litter, depending on texture and composition). We checked DNA extraction quality and concentration in a Qubit 30® fluorimeter (Invitrogen, Sweden). The soil and litter samples from which DNA was successfully extracted were sent to Aimethods (Germany) for amplification and sequencing. We targeted prokaryotes with the V3-V4 region (∼460 bases) of the 16S rDNA gene using the forward primer (5’-CCTACGGGN GGCWGCAG-3’) and the reverse primer (5’-GACTACH VGGGTATCTAATCC-3’) from Klindworth et al. (2013). Eukaryotes were targeted with the V7 region of the 18S rDNA gene using the forward and reverse primers (5’-TTTGTCTGSTTAATTSCG-3’) and (5’-TCACAGACCTGTTATTGC-3’) designed by Guardiola et al. (2015) to yield 100–110 bases long fragments. The 16S rDNA fragment was sequenced with the Illumina MiSeq 2×300 platform, and the 18S rDNA fragment with Illumina Microarray 2×150. We sequenced negative controls in all steps: three for the extraction, two for the amplification, and two for the index ligation.

### 2.6. Sequence analyses and taxonomic assessment

We used the Cutadapt package (Martin, 2011) in Python v.3.3 (Van Rossum and Drake, 2009) to remove primers. We then used the DADA2 package (Callahan et al., 2016) in R v. 4.0.2 (R Core Team, 2020) to quality filter reads, merge sequences, remove chimeras, and to infer amplicon sequence variants (ASVs). We excluded reads with ambiguous bases (maxN=0). Based on the quality scores of the forward and reverse sequences, each read was required to have <3 or <5 errors, respectively (maxEE=c (3,5), truncQ=2). Therefore, ASVs were inferred for forward and reverse reads for each sample using the run-specific error rates. To assemble paired-end reads, we considered a minimum of 12 base pairs of overlap and excluded reads with mismatches in the overlapping region. Chimeras were removed using the consensus method of “removeBimeraDenovo” implemented in DADA2. We removed ASVs present in negative controls in a proportion larger than 40% of the reads for 18S and all ASVs present in negative control for 16S. We used the SILVAngs 132.1 reference database (Quast et al., 2012) for assessment of the taxonomic composition of the ASVs for both markers. The ASV reads by sample and taxonomic affiliation are provided in the Appendix 1 (for 16S) and Appendix 2 (for 18S). Additionally, we identified the functional guild for the ASVs assigned to the fungal kingdom using the FungalTraits database (Polme et al., 2020).

### 2.7. Statistical analysis

We conducted all analyses in R using RStudio (2015). We used the tidyverse package v. 1.3.0 (Wickham, 2017) for data curation and ggplot2 v. 3.3.2 (Wickham, 2016), ggfortify v. 0.4.11 (Tang et al., 2016), gridExtra v. 2.3 (Auguie and Antonov, 2016), and ggpubr v. 0.4.0 (Kassambara and Kassambara, 2020) for data visualisation (scripts in Appendix 3).

#### 2.7.1. Soil properties

To compare our results with other areas, we included the soil property data from terra firme and várzea in Benjamin Constant (far western Brazilian Amazonia) and Caxiuanã (far eastern Amazonia), available in Ritter et al. (2018), in our data analyses (Appendix 4 Table A1). We first normalized all soil variables to zero mean and unit variance using the “scale’” function of vegan v. 2.4-3 (Oksanen et al., 2010). We then performed a principal component analysis (PCA) to reduce the number of soil property variables for subsequent analyses and visualise soil physicochemical properties in relation to forest type and flood level (i.e. terra firme, high-várzea, mid-várzea, low-várzea, or várzea where information on placement along the flooding gradient was absent).

#### 2.7.2. Alpha diversity

As the richness estimates could be biased by rare ASVs (Haegeman et al., 2013), we calculated ASV Fisher’s alpha diversity (i.e., the relationship between the number of ASVs in any given plot and the number of reads of each ASV) using the phyloseq R package v.1.34.0 (McMurdie and Holmes, 2013) separately for the prokaryote (16S) and eukaryote (18S) datasets. For the woody plant communities, we used an abundance species matrix. We calculated the metrics within each plot and compared visually the non-normalized Fisher’s alpha diversity indices of the below-ground biota and above-ground plant communities. We analysed soil and litter Fisher’s alpha diversity as a function of flood level (modelled as a continuous variable represented by the measured floodwater marks on trees, with terra firme being zero, and categorically according to forest type, i.e. flood level), soil properties (represented by PC1 of the soil PCA), type of sample (litter or soil), and above-ground Fisher’s alpha diversity for woody plants. We normalized all the Fisher’s alpha diversities to zero mean and unit variance using the “scale’” function in vegan. Thus, we defined a set of models to explain below-ground alpha diversity. The final model set included models with flood level, inundation depth of the last flood, PC1 from the soil properties PCA, type of sample (litter or soil) and woody plant Fisher’s alpha diversity as predictor variables, and additional models with interaction terms among the flood levels and sample types with woody plant Fisher’s alpha diversity and the flood levels with soil PC1. The final model set also included a constant, intercept-only model, comprising a total of nine models for each dependent variable (Table 1).

**Table 1.**
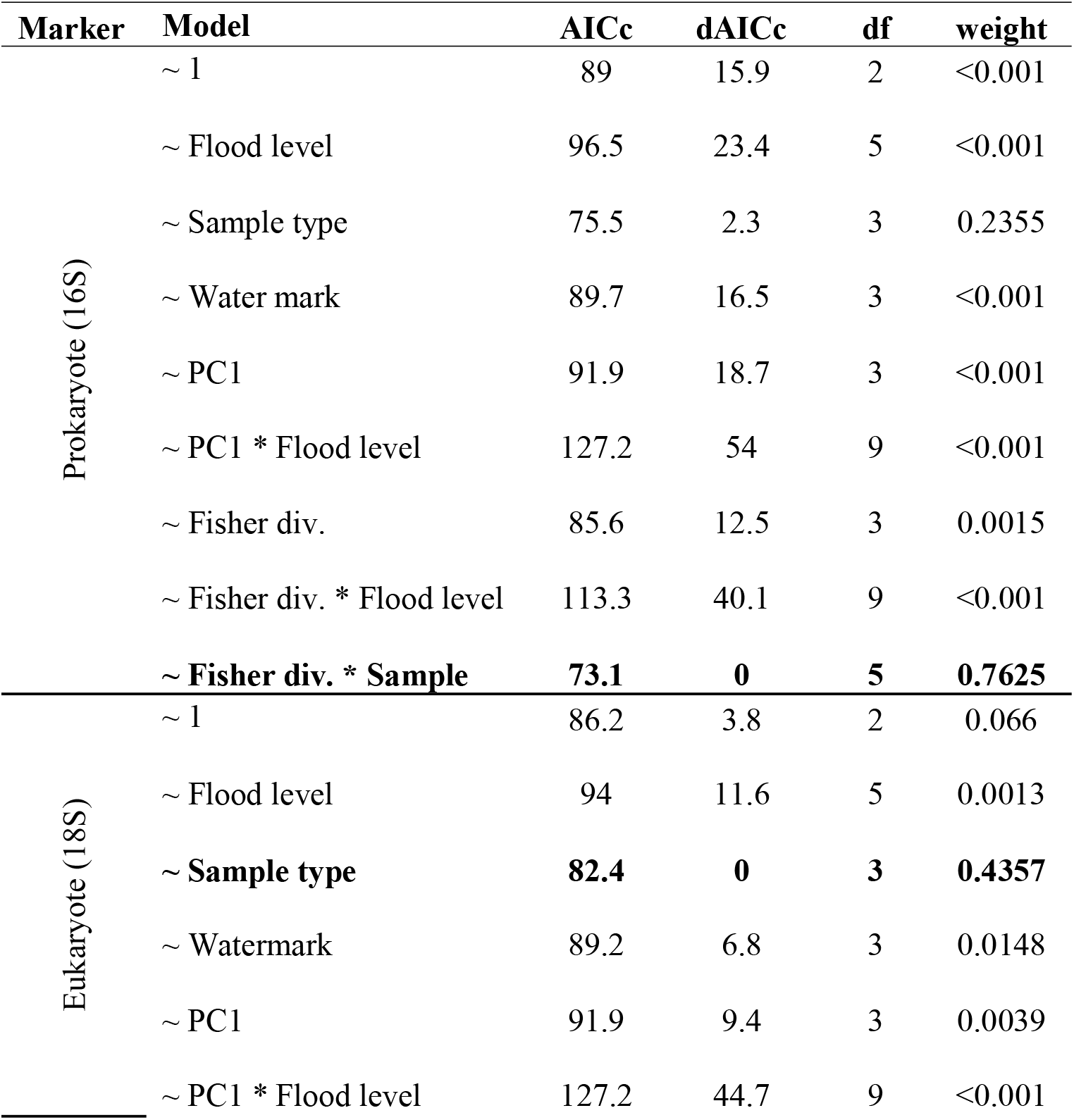
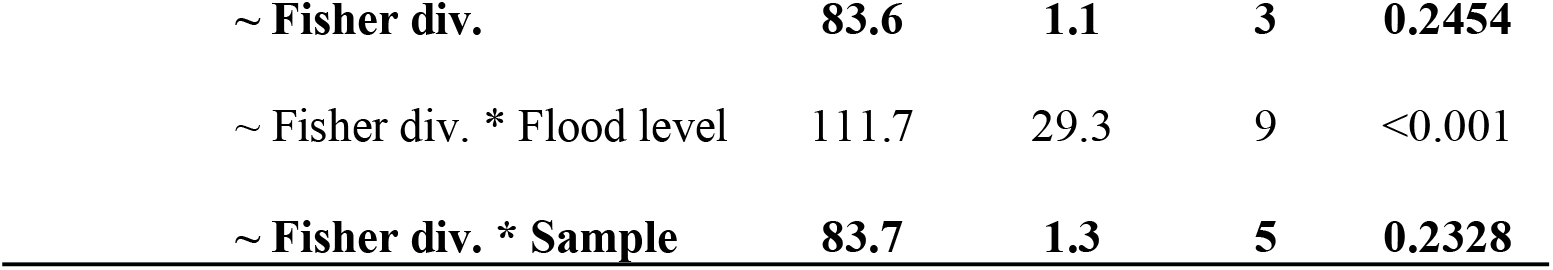
Variables used in model selection with their respective delta dAICc and weight values. The best fit model has a dAICc = 0 and is presented in bold as the alternative good models (dAICc =< 2). The response variables are below-ground Fisher’s diversity for prokaryotes (16S) and eukaryotes (18S). The independent variables are flood level, sample type, water mark (measured floodwater marks on trees, with terra firme being zero), and the woody plant Fisher’s diversity. The model used flood level and sample type as a fixed factor or as interacting variable.

| Marker | Model | AICc | dAICc | df | weight |
| --- | --- | --- | --- | --- | --- |
| Prokaryote (16S) | ~ 1 | 89 | 15.9 | 2 | <0.001 |
|  | ~ Flood level | 96.5 | 23.4 | 5 | <0.001 |
|  | ~ Sample type | 75.5 | 2.3 | 3 | 0.2355 |
|  | ~ Water mark | 89.7 | 16.5 | 3 | <0.001 |
|  | ~ PC1 | 91.9 | 18.7 | 3 | <0.001 |
|  | ~ PC1 * Flood level | 127.2 | 54 | 9 | <0.001 |
|  | ~ Fisher div. | 85.6 | 12.5 | 3 | 0.0015 |
|  | ~ Fisher div. * Flood level | 113.3 | 40.1 | 9 | <0.001 |
|  | **~ Fisher div. \* Sample** | **73.1** | **0** | **5** | **0.7625** |
| Eukaryote (18S) | ~ 1 | 86.2 | 3.8 | 2 | 0.066 |
|  | ~ Flood level | 94 | 11.6 | 5 | 0.0013 |
|  | **~ Sample type** | **82.4** | **0** | **3** | **0.4357** |
|  | ~ Watermark | 89.2 | 6.8 | 3 | 0.0148 |
|  | ~ PC1 | 91.9 | 9.4 | 3 | 0.0039 |
|  | ~ PC1 * Flood level | 127.2 | 44.7 | 9 | <0.001 |

|  |  |  |  |  |
| --- | --- | --- | --- | --- |
| ~ **Fisher div.** | **83.6** | **1.1** | **3** | **0.2454** |
| ~ Fisher div. * Flood level | 111.7 | 29.3 | 9 | <0.001 |
| ~ **Fisher div. \* Sample** | **83.7** | **1.3** | **5** | **0.2328** |

Models were selected using an information theory approach based on AIC (Akaike, 1974) and corrected AICs (AICc) for small sample sizes (Burnham and Anderson, 2002). Models with dAIC ≤ 2 were considered equally plausible, and we used the normalized model weight (wi) to contrast the best model to the constant (no-effect) model. We used generalized linear models (Crawley, 2007) with Gaussian error distributions after checking for the distributions of residuals. The GLM analyses were performed using the vegan package, and model selection was carried out using the bbmle package v.1.0.20 (Bolker and Bolker, 2017).

#### 2.7.3. Beta diversity

We constructed two-dimensional non-metric multidimensional scaling (NMDS) ordinations of the abundance (reads) matrices of prokaryotes (16S) and eukaryotes (18S). We first transformed read counts using the ‘varianceStabilizingTransformation’ function in DESeq2 v.1.30.1 (Love et al., 2014) as suggested by McMurdie & Holmes (2013). This transformation normalizes the count data with respect to sample size (number of reads in each sample) and variances, based on fitted dispersion-mean relationships (Love et al., 2014). We then used the ‘metaMDS’ function and Bray-Curtis distances in the vegan package to assess community dissimilarity among all samples in the NMDS. We used the ‘envfit’ method in vegan to fit flood levels and sample types onto the NMDS ordination as a measure of the correlation among these factors with the NMDS axes. Additionally, we constructed two-dimensional non-metric multidimensional scaling (NMDS) ordinations based on the abundance data of the woody plants.

## 3. Results

We were able to extract, amplify, and sequence DNA for both prokaryotes (16S) and eukaryotes (18S) in 13 soil samples, 17 litter samples for prokaryotes (16S), and 16 litter samples for eukaryotes (18S). We obtained a total of 787,834 reads and 10,213 ASVs for the prokaryotes (16S). After removing the negative controls, we kept 757,827 reads and 9,337 ASVs. For the eukaryotes (18S), we obtained 616,237 reads belonging to 2,267 ASVs and we kept 572,953 reads belonging to 2,004 ASVs after removing the negative controls. See Appendix 4, Table A2 for the number of reads and ASV richness for each plot, and Appendix 5 and 6 for krona charts of 16S and 18S taxonomic composition, respectively. The raw sequences are deposited in Genbank under the Bioproject PRJNA723037, BioSample SAMN18800640: Jurua (TaxID: 410658), accession SRA numbers: SRR14286278 - SRR14286277.

### 3.1. Soil properties

The principal component analysis showed that edaphic properties varied between terra firme and várzea plots and that flood depth or duration had no apparent effect on várzea soil physicochemical composition (Fig. 2; Appendix 4 Table A3). Hence, várzea soils from Juruá largely overlapped (Fig. 2). Várzea soils were dominated by clay and silt, whereas terra firme soils were sandier (Fig 2). Terra firme soils were less fertile than várzea soils, with lower concentrations of important nutrients, such as potassium (K), calcium (Ca), and magnesium (Mg) (Fig 2). Compared with the terra firme and várzea soils from Benjamin Constant (far western Brazilian Amazonia) and Caxiuanã (far eastern Brazilian Amazonia), the Juruá várzea is characterized by more exchangeable bases and clay, and less phosphorous (P). The Juruá terra firme soils are placed between the Benjamin Constant and Caxiuanã terra firme soils (Fig. 2).

**Fig. 2.**
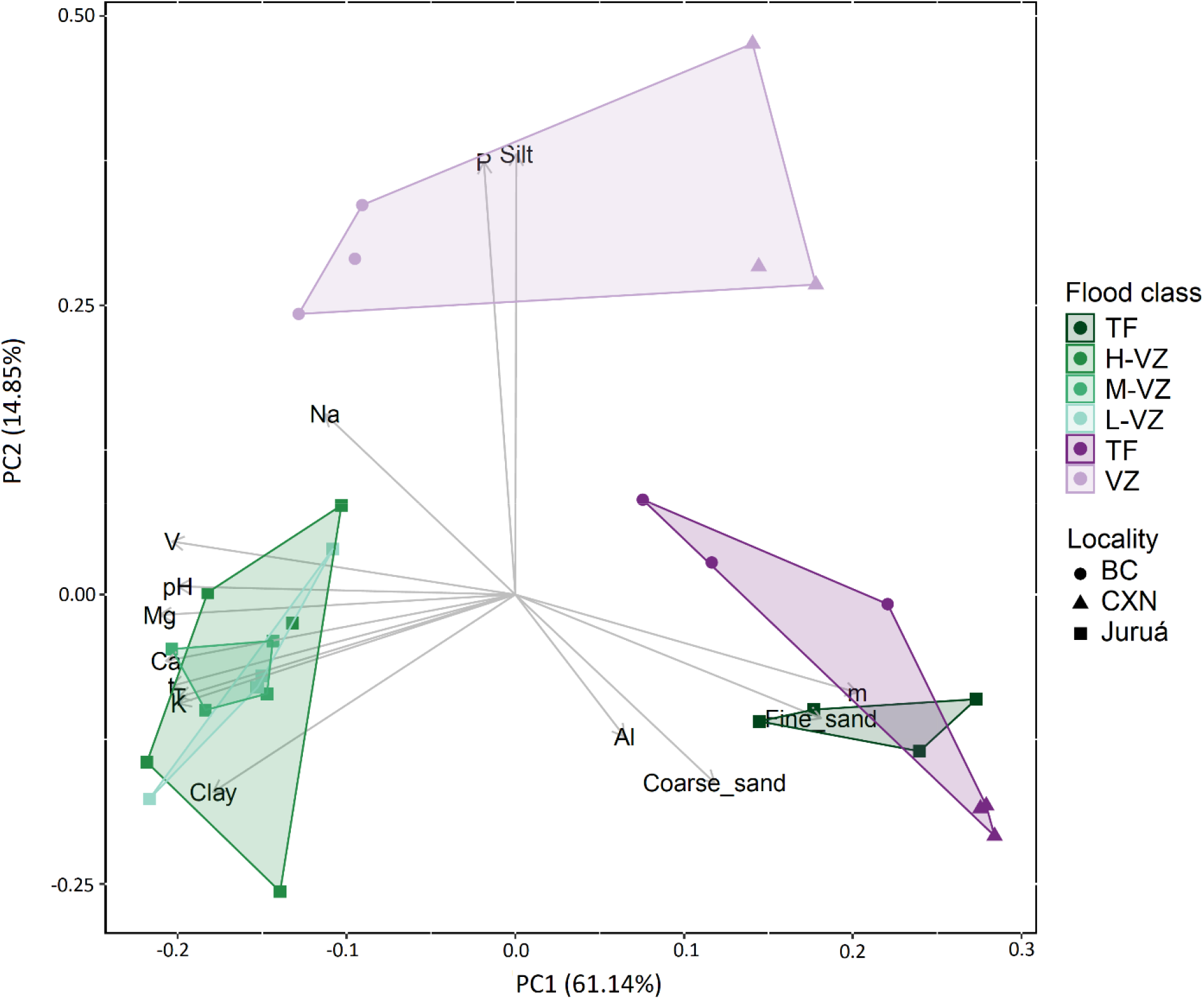
Principal component analysis (PCA) showing the clustering of inventory plots along the first two PCA axes in relation to the soil physicochemical composition. The colours of the clusters reveal the geographic location (Juruá - this study - in green nuances; Benjamin Constant and Caxiuanã = purple) and the flooding gradient represented by the Juruá flood levels: TF: Terra firme; HV: High-várzea; MV: Mid-várzea; and LV: Low-várzea. The shape of the points indicates plot locality: Juruá = squares, Benjamin Constant = circle; and Caxiuanã = triangles.

### 3.2. Below-ground taxonomic composition

The taxonomic composition of the prokaryote component shows that the groups with the highest number of ASVs were Alphaproteobacteria (∼25% of the taxa identified in our samples, equivalent to ∼2000 ASVs per flood level; Fig. 3A; Appendix 4 Fig. A1A), Actinobacteria (∼23%, average ∼1700 ASVs; Fig. 3A; Appendix 4 Fig. A1A), and Acidobacteria (∼18%, average ∼1300 ASVs; Fig. 3A; Appendix 4 Fig. A1A). Among eukaryotes, Fungi had the highest number of ASVs (∼43%, ∼600 ASVs), mainly Ascomycota and Basidiomycota (Fig. 3B; Appendix 4 Fig. A1B) followed by Cercozoa (∼18%, ∼300 ASVs; Fig. 3B; Appendix 4 Fig S1B) and Ciliophora (∼15%, ∼250 ASVs; Fig. 3B; Appendix 4 Fig A1B). Most fungi were classified as saprotrophs (Appendix 4 Fig. A2). Other groups present were pathogens, parasites, mycorrhizae fungi and unclassified (Appendix 4 Fig. A2).

**Fig. 3:**
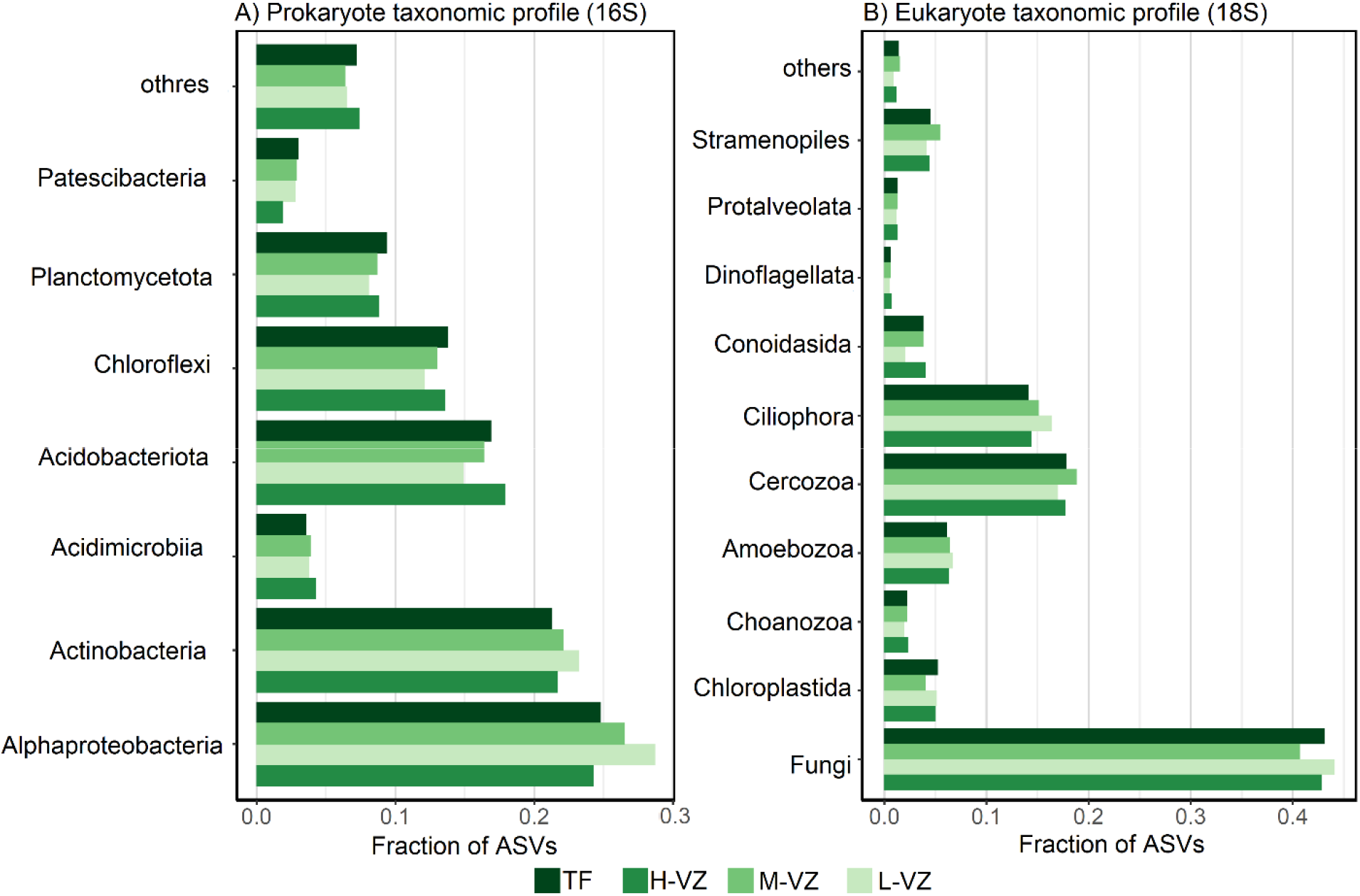
Fraction of ASVs by taxonomic group and flood level for (A) prokaryotes and (B) eukaryotes. Flood levels are TF: Terra firme; H-VZ: High-várzea; M-VZ: Mid-várzea; and L-VZ: Low-várzea.

### 3.3. Alpha diversity

We found that the best model to explain bacterial (16S) diversity included woody plant Fisher’s alpha diversity and sample type (soil or litter) with an interaction effect between the two (Table 1), but only sample type was significant (Table 2). For eukaryotes (18S), three models had a delta AICc lower than 2 (Table 1). The first model (dAICc = 0) included only sample type, the second (dAICc = 1.1) included only the woody plant Fisher’s alpha diversity, and the third model (dAICc = 1.3) included the woody plant Fisher’s alpha diversity and sample type with an interaction effect between the two (Table 1). In all models, only sample type was significant (Table 2). Bacterial Fisher’s alpha diversity was higher than the Fisher’s alpha diversity of either eukaryotes or woody plants. In terra firme, bacterial diversity in soil and litter, but not eukaryotes, appears to correlate with woody plant diversity. For várzea, no pattern was observed (Fig. 4).

**Table 2.** Estimated parameters (values estimated with standardize error, t-value and respective p-value) of the best fit model for 16S and the third best fit model (that included the variables selected) for 18S selected in model selection. The response variables are below-ground Fisher’s alpha diversity and (above-ground) woody plant Fisher’s alpha diversity with an interaction term between the above-ground alpha diversity and sample type (soil or litter). Significant factors (p < 0.05) are marked in bold.

|  | Coefficients | Estimate | Std.<br>Error | t value | Pr(>\|t\|) |
| --- | --- | --- | --- | --- | --- |
| Prokaryote (16S) | (Intercept) | 0.6427 | 0.1815 | 3.54 | 0.00167 |
|  | fisher.alpha | -0.2288 | 0.1947 | -1.175 | 0.2515 |
|  | **SampleSoil** | **-1.3463** | **0.2774** | **-4.853** | **6.03E-05** |
|  | fisher.alpha:SampleSoil | 0.4439 | 0.2797 | 1.587 | 0.12566 |
| Eukaryote (18S) | (Intercept) | 0.43232 | 0.23733 | 1.822 | 0.0816 |
|  | fisher.alpha | -0.03453 | 0.25159 | -0.137 | 0.892 |
|  | **SampleSoil** | **-0.89964** | **0.35607** | **-2.527** | **0.0189** |
|  | fisher.alpha:SampleSoil | 0.43738 | 0.36073 | 1.212 | 0.2376 |

**Fig. 4.**
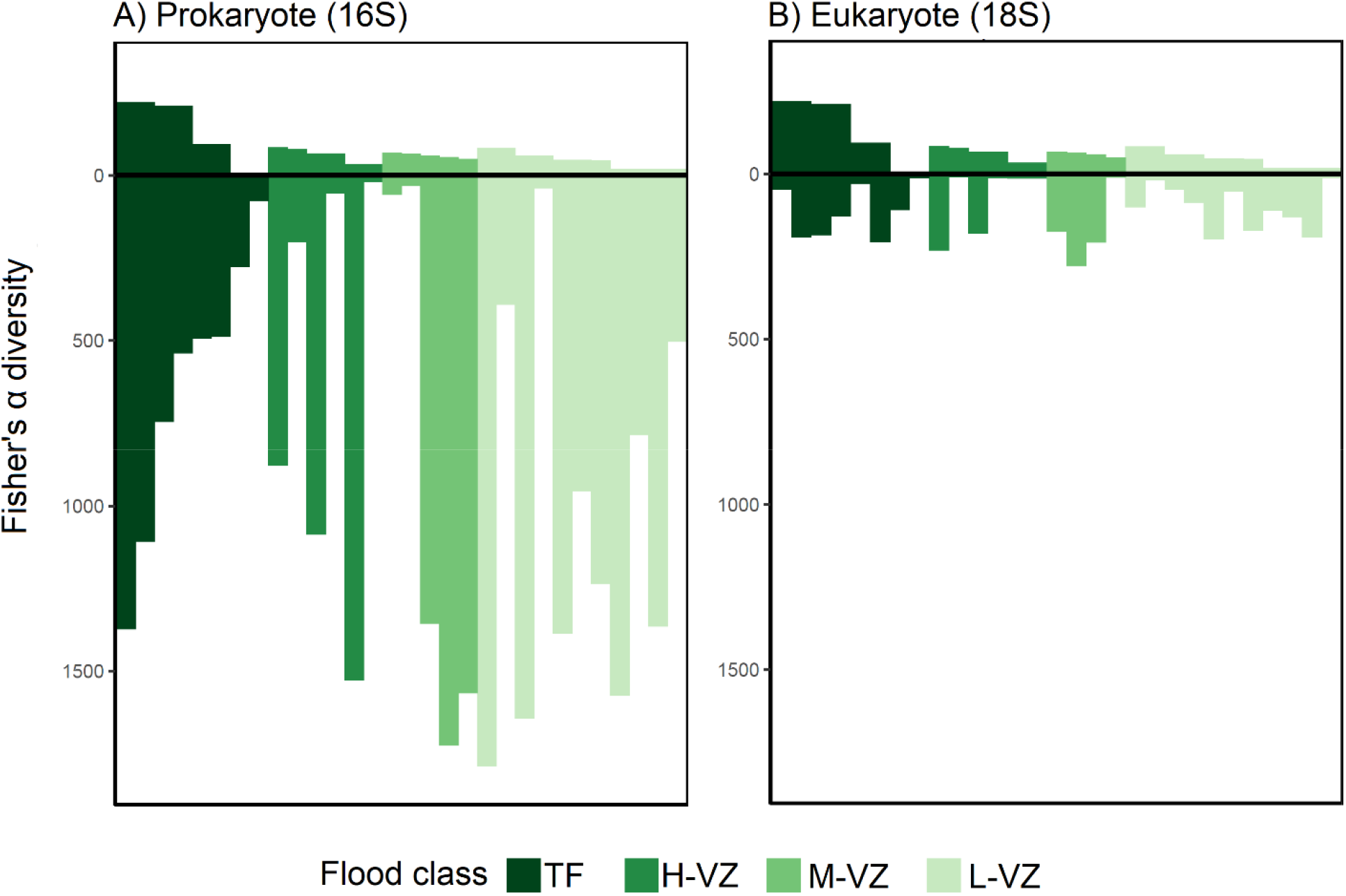
Above-ground woody plant Fisher’s alpha diversity *versus* below-ground Fisher’s alpha diversity of A) prokaryotic (16S) and B) eukaryotic (18S) organisms in Juruá litter and soil samples. Prokaryotic and eukaryotic diversity are shown in negative values. Woody plant diversity is shown in positive values. Flood levels are TF: Terra firme; H-VZ: High-várzea; M-VZ: Mid-várzea; and L-VZ: Low-várzea.

### 3.4. Beta diversity

Community compositions were similar among plots across flood levels and sample types (litter and soil). For bacteria, there is a grouping of terra firme plots with some overlap with várzea plots (Fig. 5A). No clear pattern was observed for soil eukaryotes (Fig. 5B). For woody plant communities, there is a turnover in species compositions across different flood levels (Fig. 5C). The envfit test indicated a significant effect of flood level on both the prokaryote (R^2^ = 0.24; p = 0.022) and woody plant (R^2^ = 0.48; p = 0.003) communities, but not for soil eukaryotes (R^2^ = 0.14; p = 0.28). The envfit test also indicated a significant effect of sample type on the prokaryote (R^2^ = 0.25; p = 0.001) and eukaryote (R^2^ = 0.22; p = 0.006) communities.

**Fig. 5.**
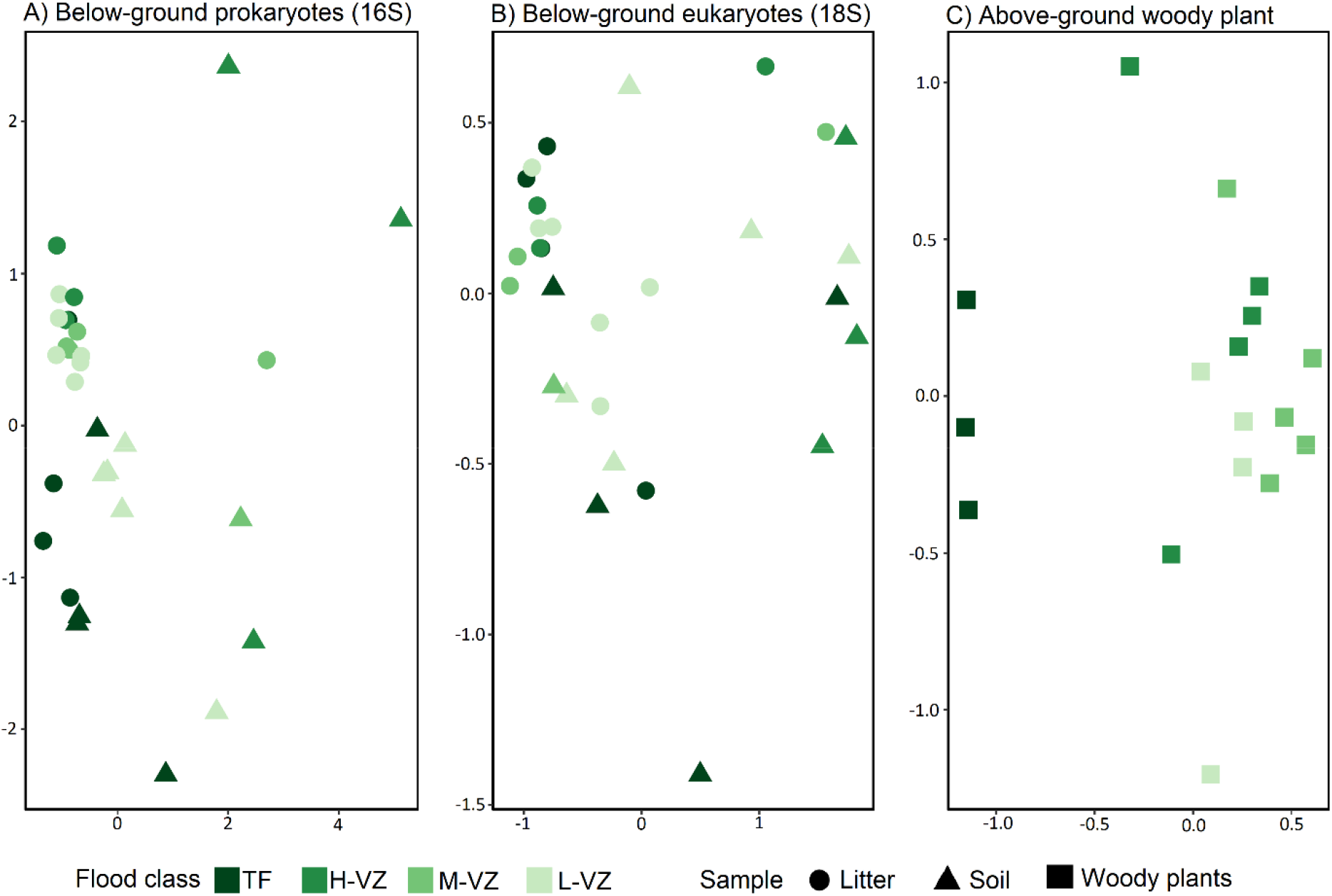
Community structure in relation to substrate type and flood levels. Visualisation of non-metric multidimensional scaling (NMDS) for (A) prokaryotes (16S), (B) eukaryotes (18S), and (C) woody plants using Bray-Curtis dissimilarity indices. Symbols represent different substrates (i.e. sample types) where filled circles = litter samples and filled triangles = soil samples. Colours represent the different flood levels: TF = Terra firme; H-VZ = High-várzea; M-VZ = Mid-várzea; and L-VZ = Low-várzea.

## 4. Discussion

Our analyses have documented, for the first time, the degree to which soil and litter biota biodiversity are affected by the flooding gradient in central-western Amazonian forests of varying floristic diversity. We show a weak correlation between soil and litter community composition and inundation period but find that below-ground Fisher’s alpha diversity cannot be explained by the flooding gradient. We also show that the edaphic properties differed between terra firme and várzea, but not among várzea forests along the flooding gradient.

### 4.1. Edaphic properties

Várzea edaphic properties in the Juruá differed from the other two Amazonian várzeas that we included in our analyses (Fig. 2). For instance, the Juruá várzea was poorer in phosphorus (P) and silt, but rich in magnesium (Mg), calcium (Ca), potassium (K), and clay. This high-density clay content in the Juruá várzea may act as a physical barrier to water infiltration. On the other hand, clayey soils also have a high water holding capacity (Hillel, 2013), which prevents it from drying out completely during the non-flooded periods. The high clay content additionally made várzea samples hard to collect and to break once dried. Possibly, this was the main factor that hindered DNA extraction in our study.

Compared to the terra firme soils, the Juruá várzea soils were more fertile, presumably due to the yearly inflow of nutrient-rich alluvial sediments by the Juruá River. Moreover, the Juruá terra firme soils presented similar edaphic properties to those of the terra firme forests in Benjamin Constant and Caxiuanã. This was unexpected since the terra firme forest that we sampled in the Juruá grow on paleo-várzea sediments (Assis et al., 2015), and therefore presumably should have been relatively nutrient rich compared to typically well-drained and heavily leached terra firme soils on older geological formations (Sombroek, 2000). However, these soils presented similar edaphic properties to those of the terra firme forests in Benjamin Constant and Caxiuanã, suggesting that nutrients are soon leached from várzea substrates once they no longer experience flooding and an influx of river sediments.

### 4.2. Below-ground taxonomic composition

Alphaproteobacteria and Planctomycetes were abundant in our samples, accounting for 40% of our 16S data (Fig. 3A). These groups are known to be very diverse in undisturbed forests (de Carvalho et al., 2016) and they are generally common in Amazonian soils (Ritter et al., 2019b; Zinger et al., 2019). Interestingly, other bacterial groups commonly found in Amazonian soils (Ritter et al., 2019b; Zinger et al., 2019) and elsewhere (Delgado-Baquerizo et al., 2018) – notably Betaproteobacteria and Bacteroidetes - were not present in the Juruá samples. Because these groups are known form a diverse range of habitats, including várzea and terra firme, this surprised us and clearly highlight that we have much to discover about Amazonian soil biodiversity.

Patescibacteria (e.g., the candidate phyla radiation group), not previously reported in other várzea soils, were found in the Juruá samples (Fig. 3A). This group was recently described (Brown et al., 2015) and until now it had only been registered in Amazonian pasture soils (Lemos et al., 2020). An interesting characteristic of Patescibacteria is the small size of their genomes (usually <1.5 Mbp) and their lack of biosynthetic capabilities (Brown et al., 2015). These characteristics indicate that they could be co-metabolic interdependent (He et al., 2015; Lemos et al., 2019). Such interdependencies with other organisms would suggest a restrict occurrence or different functionality dependent on the community in which they occur. Yet, Patescibacteria show similar functional profiles under distinct climate conditions (tropical soils and permafrost; Lemos et al., 2020). Although their apparent plasticity is interesting, very little information is available for this group. The design of new 16S rRNA gene primers that better amplify Patescibacteria is required to elucidate the ecology and distribution of Patescibacteria in Amazonian soils and worldwide. Additionally, analysis of metatranscriptomes could improve our understanding of the metabolism in Patescibacteria and other bacteria under different substrate conditions.

Among the eukaryotes, we found a higher proportion of fungi in the Juruá substrates than previously documented for other areas in Amazonia (Ritter et al., 2019b). Whereas Ritter et al. (2020) found fewer fungi in várzea than in other environments, we found more fungi in várzea than in the adjacent terra firme, most of which were saprotrophs (Appendix 4 Fig. A2). Singer et al. (1983) hypothesized that ectomycorrhizal fungi increase the ability of their host plants to acquire nutrients and water in low-fertility soils, such as in the Amazonian sandy-soil ecosystems. However, we found very few ectomycorrhizal fungi in both várzea (more fertile) and terra firme (less fertile; Appendix 4 Fig. A2). Yet, around 35% of the fungi could be not assigned to any functional guild. This makes comparisons difficult and highlights the need to further investigate Amazonian soil biodiversity and its ecology.

Some eukaryotic groups detected in other Amazonian localities by the same 18S primers as the ones used here (Ritter et al., 2019b; Zinger et al., 2019), were absent in the Juruá samples. Such groups include nematodes and arthropods (Fig. 2B). Although the 18S primers that we used are not optimal for sequencing animals, it was surprising not to find these groups in our samples (except for one nematode sequence in várzea and terra firme). Low nematode diversity in Amazonian várzeas was previously reported by Cares (1984). One reason for the absence of these animals in várzea substrates could be that the high amount of clay in the soil and the seasonal floods, make várzea soil and litter unfit for nematode occupation. However, this does not explain the absence of soil animals in our terra firme samples since these were relatively clay-free and unflooded. To test this hypothesis, we need further studies in soils with a gradual difference in clay proportion and specific primers targeting nematodes (e.g. Kawanobe et al., 2021) alongside morphological examination of the diversity in the samples.

### 4.3. Above-versus below-ground diversity

There was no relationship between above- and below-ground alpha diversity across the different forest types included in this study. This mismatch could be explained by the flood pulse that may have masked any pattern by carrying organisms across all flood levels. A lack of clear relationships between above- and below-ground biodiversity has previously been demonstrated globally (Cameron et al., 2019) and for other Amazonian areas (Ritter et al., 2019a). However, for Amazonia this mismatch was partial. Across habitats, no correspondence was found between below-ground prokaryote or eukaryote alpha diversity and above-ground bird or tree alpha diversity (Ritter et al., 2019a). Nevertheless, there was a gradual decrease in below- and above-ground alpha diversity from the west to the east across the Amazon basin (Ritter et al., 2019a). Indeed, bacterial diversity appears to correlate with woody plant diversity in terra firme forests (Fig. 3A), but due to the sample limitation, just four terra firme plots, we could not find a significance in this relationship.

### 4.4 Flooding gradient and community composition

Most ASVs occur throughout the flooding gradient (Appendix 1 and 2). This result was partly expected since the seasonal flood waters could carry DNA (e.g. of inactive spores, dead or living organisms) across all várzea flood levels. Yet, the bacterial community composition of the Juruá substrate varied with flood level and woody plant diversity. This result indicates that below-ground bacteria may present different tolerances to hydrological stressors and or interdependencies with certain woody plant species. For instance, nodulation caused by nitrogen fixing bacteria are more frequent in Amazonian seasonally flooded forests, indicating that nodulation may be favored in flooded areas (Parolin and Wittmann, 2010).

## 5. Conclusion

This is the first study to investigate the degree to which soil and litter biota are affected by the flooding gradient in Amazonian forests. In fact, as far as we are aware, substrates from only six other Amazonian várzeas have previously been investigated using a metabarcoding approach, and these studies did not consider the flooding gradient (Ritter et al., 2019b, 2019a; Ritter, 2018). Hence, the DNA barcoding data herein – consisting of a total of 19,550 ASVs, from 14 várzea and four terra firme plots – more than doubles the total database from Amazonian várzeas available to date. Considering the extent of lowland Amazonian floodplain forests, approx. 516,000 km^2^ (Hess et al., 2015) the need for more data from different geographical areas is obvious.

Studying below-ground communities along complex environmental gradients, like the one in the present study, offers an excellent opportunity to explore the responses of substrate biota to varying degrees of environmental stressors. Such studies can further our understanding of the patterns in below-ground biodiversity, their roles in the dynamics of seasonally flooded forests, and how these communities might respond to anthropogenic pressure and climate change. Therefore, the characterization of below-ground biodiversity in flooded forests, has theoretical implications for elucidating the patterns of biological diversity distribution. Practical implications include the identification of strategically important areas or areas of greater environmental sensitivity, for the conservation of biological diversity in face of environmental change. This not trivial, as infrastructural development (e.g. hydroelectric dams) and climate change (more frequent extreme floods and droughts) are severely affecting the natural flood pulse and threaten the ecological integrity of seasonally flooded forests across Amazonia (Gloor et al., 2013; Junk et al., 2018; Latrubesse et al., 2020). Increased pressures in these ecosystems highlight the urgency for more studies of this kind to improve our understanding of biodiversity patterns and community structures as these will allow us to better foresee and mitigate climate change impacts on ecosystem functions.

## Acknowledgements

This publication is part of the Projeto Médio Juruá series on Resource Management in Amazonian Reserves (www.projetomediojurua.org). We thank the Secretaria do Estado do Meio Ambiente e Desenvolvimento Sustentável do Amazonas (SEMA - DEMUC) and Instituto Brasileiro do Meio Ambiente e Recursos Naturais Renováveis (IBAMA)/Instituto Chico Mendes de Conservação da Biodiversidade (ICMBio) for authorising the research nº60/2016 (RDS Uacari) and nº55077-1 (RESEX Médio Juruá). We are grateful to the members from the Projeto Médio Juruá team for logistical support. We thank Rita Homem Pelicano for dedicated soil sampling in the field! We thank Michael J.G. Hopkins, Rafael Leandro de Assis and Juliana Schietti with colleagues at INPA for providing logistical assistance and a place to work in Manaus. We thank Paulo Apóstolo Costa Lima Assunção (in memoriam) and Alexandro Elias dos Santos, assisted by Lorena M. Rincón, for identifying the woody plant species in the field and at the National Institute for Amazonia Research (INPA). We thank our local field assistants, Associação dos Produtores Rurais de Carauari (ASPROC), Operação Amazônia Nativa (OPAN), and the people of the central Juruá who in various ways assisted us throughout our work. Finally, we thank Heléne Aronson and Katarzyna Wojnicka at the University of Gothenburg for quantifying the sampled DNA.

