## Appendix4 for "Above- and below-ground biodiversity responses to the prolonged flood pulse in central-western Amazonia, Brazil"

**Appendix 4**

Table A1. Physicochemical soil properties included in our analyses. Locality is the geographic region from where the sample was taken. Juruá indicates samples that we collected for the present study from the central Juruá. BC indicates samples collected in Benjamin Constante, see Ritter et al. (2018), CXN indicates samples collected in Caixuanã, see Ritter et al. (2018). Flood level indicates flood level where this information was available otherwise forest type, where TF = terra firme (not seasonally flooded); VZ = várzea (seasonally flooded); HV = high várzea (flooded 0-1 mo/yr); MV = mid várzea (flooded 2-4 mo/yr); and LV = low várzea (flooded 5-12 mo/yr). Coarse, Fine, and Total sand indicate the content of sand with grainsize 2.0-0.2 mm, 0.2-0.05 mm or 2.0-0.05 mm, respectively, in g/kg. Silt and Clay indicate clay and silt content in g/kg. pH is that of sample when dissolved in water. P = phosphorus, K = potassium, and Na = Natrium are given in mg/dm^3^. Ca = Calcium, Mg = magnesium, Al = aluminium, H+Al = exchangeable aluminium, SB = sum of exchangeable bases, t = effective cation exchange capacity, T = cation exchange capacity, V = Base Saturation Index, and m = aluminium saturation index are given in cmol_c_/dm^3^.

| Locality | Flood level | Coarse sand | Fine sand | Total sand | Silt | Clay | pH | P | K | Na | Ca | Mg | Al | H+Al | SB | t | T | V | m |
| --- | --- | --- | --- | --- | --- | --- | --- | --- | --- | --- | --- | --- | --- | --- | --- | --- | --- | --- | --- |
| Juruá | TF | 13.7 | 418.5 | 432.2 | 426.3 | 141.5 | 4.0 | 5.0 | 46.0 | 5.0 | 0.1 | 0.1 | 3.6 | 7.8 | 0.4 | 3.9 | 8.1 | 4.3 | 91.1 |
| Juruá | TF | 39.4 | 129.3 | 168.7 | 382.8 | 448.5 | 4.0 | 3.0 | 86.0 | 45.0 | 0.4 | 0.4 | 8.2 | 11.7 | 1.2 | 9.4 | 12.9 | 9.6 | 86.9 |
| Juruá | LV | 5.9 | 12.9 | 18.9 | 512.7 | 468.5 | 4.7 | 10.8 | 189.0 | 28.0 | 9.1 | 2.8 | 2.1 | 8.0 | 12.6 | 14.7 | 20.6 | 61.0 | 14.3 |
| Juruá | LV | 11.3 | 15.0 | 26.3 | 275.7 | 698.0 | 4.7 | 7.4 | 146.0 | 42.0 | 10.6 | 3.6 | 1.9 | 10.8 | 14.8 | 16.6 | 25.5 | 57.9 | 11.1 |
| Juruá | MV | 17.4 | 15.9 | 33.3 | 314.2 | 652.5 | 4.6 | 6.3 | 142.0 | 34.0 | 11.8 | 3.7 | 1.9 | 9.8 | 16.0 | 17.9 | 25.7 | 62.0 | 10.8 |
| Juruá | MV | 4.0 | 9.3 | 13.3 | 441.2 | 545.5 | 4.6 | 8.5 | 145.0 | 35.0 | 13.9 | 3.1 | 3.8 | 9.3 | 17.5 | 21.4 | 26.9 | 65.2 | 17.9 |
| Juruá | MV | 11.8 | 15.6 | 27.3 | 337.7 | 635.0 | 4.6 | 8.4 | 147.0 | 22.0 | 14.8 | 3.2 | 2.0 | 2.4 | 18.4 | 20.4 | 20.8 | 88.7 | 9.7 |
| Juruá | HV | 7.1 | 7.9 | 15.0 | 565.0 | 420.0 | 4.8 | 11.3 | 139.0 | 23.0 | 9.7 | 3.2 | 1.7 | 7.7 | 13.4 | 15.1 | 21.1 | 63.4 | 11.4 |
| Juruá | HV | 11.5 | 32.4 | 43.9 | 6.6 | 949.5 | 4.6 | 11.0 | 164.0 | 20.0 | 11.7 | 2.9 | 3.2 | 9.0 | 15.1 | 18.2 | 24.1 | 62.6 | 17.3 |
| Juruá | HV | 10.1 | 10.3 | 20.4 | 533.6 | 446.0 | 4.6 | 8.1 | 156.0 | 23.0 | 12.3 | 4.0 | 3.5 | 9.5 | 16.8 | 20.3 | 26.3 | 63.9 | 17.3 |
| Juruá | HV | 7.1 | 16.3 | 23.4 | 282.1 | 694.5 | 5.0 | 7.0 | 172.0 | 23.0 | 14.8 | 4.8 | 0.8 | 14.2 | 20.1 | 20.9 | 34.3 | 58.7 | 4.0 |
| Juruá | MV | 16.5 | 14.3 | 30.7 | 295.8 | 673.5 | 5.1 | 4.8 | 158.0 | 28.0 | 12.6 | 3.8 | 0.5 | 7.4 | 16.9 | 17.4 | 24.3 | 69.7 | 2.6 |
| Juruá | TF | 12.2 | 13.2 | 25.4 | 416.6 | 558.0 | 3.8 | 6.2 | 69.0 | 5.0 | 0.2 | 0.2 | 4.2 | 11.7 | 0.5 | 4.8 | 12.3 | 4.5 | 88.5 |
| Juruá | MV | 8.0 | 10.1 | 18.1 | 359.4 | 622.5 | 4.9 | 8.1 | 167.0 | 34.0 | 14.7 | 4.0 | 0.8 | 8.6 | 19.3 | 20.0 | 27.9 | 69.0 | 3.7 |
| Juruá | LV | 6.8 | 12.1 | 18.9 | 321.6 | 659.5 | 4.8 | 7.3 | 156.0 | 26.0 | 12.2 | 3.2 | 1.1 | 8.9 | 16.0 | 17.0 | 24.8 | 64.3 | 6.2 |
| Juruá | LV | 3.0 | 13.7 | 16.7 | 344.3 | 639.0 | 4.4 | 7.0 | 161.0 | 28.0 | 20.9 | 4.1 | 4.1 | 12.6 | 25.6 | 29.6 | 38.1 | 67.1 | 13.7 |
| Juruá | HV | 2.1 | 8.2 | 10.3 | 418.2 | 571.5 | 4.7 | 11.1 | 179.0 | 36.0 | 14.6 | 3.3 | 1.5 | 8.6 | 18.5 | 19.9 | 27.1 | 68.3 | 7.3 |
| Juruá | TF | 54.7 | 296.8 | 351.5 | 395.1 | 253.5 | 3.7 | 8.0 | 81.0 | 6.0 | 0.1 | 0.3 | 6.0 | 13.1 | 0.6 | 6.6 | 13.8 | 4.6 | 90.4 |
| BC | TF | 10.8 | 127.4 | 138.2 | 524.6 | 337.2 | 4.3 | 10.3 | 86.0 | 18.7 | 3.1 | 1.2 | 1.9 | 6.2 | 4.7 | 6.6 | 10.9 | 42.3 | 30.1 |
| BC | TF | 9.5 | 110.9 | 120.4 | 527.6 | 352.0 | 4.1 | 10.3 | 64.0 | 12.0 | 2.5 | 1.4 | 4.2 | 7.1 | 4.0 | 8.3 | 11.1 | 36.1 | 51.1 |
| BC | TF | 9.1 | 329.2 | 338.3 | 414.6 | 247.2 | 3.9 | 9.3 | 49.7 | 25.7 | 0.5 | 0.3 | 4.3 | 7.3 | 1.0 | 5.3 | 8.3 | 12.2 | 80.9 |
| BC | VZ | 1.4 | 4.3 | 5.8 | 591.1 | 403.2 | 4.5 | 32.0 | 94.3 | 36.0 | 9.8 | 3.1 | 1.5 | 4.2 | 13.3 | 14.8 | 17.5 | 75.8 | 10.3 |
| BC | VZ | 1.3 | 2.7 | 4.0 | 568.3 | 427.7 | 4.7 | 29.5 | 96.0 | 29.7 | 10.6 | 2.8 | 1.3 | 3.7 | 13.7 | 15.0 | 17.4 | 78.8 | 8.5 |
| BC | VZ | 5.4 | 4.7 | 10.2 | 521.0 | 468.8 | 4.6 | 23.3 | 97.0 | 48.7 | 11.4 | 3.5 | 1.6 | 5.4 | 15.3 | 16.9 | 20.7 | 73.9 | 9.4 |
| CXN | TF | 531.5 | 272.7 | 804.2 | 62.3 | 133.5 | 3.9 | 5.1 | 22.7 | 14.0 | 0.2 | 0.2 | 1.2 | 4.6 | 0.5 | 1.6 | 5.1 | 9.1 | 72.0 |
| CXN | TF | 488.0 | 269.7 | 757.7 | 81.6 | 160.7 | 3.7 | 5.9 | 27.0 | 21.3 | 0.1 | 0.2 | 1.5 | 5.7 | 0.4 | 1.9 | 6.1 | 7.2 | 77.2 |
| CXN | TF | 557.0 | 234.2 | 791.2 | 59.8 | 149.0 | 3.8 | 8.2 | 36.0 | 18.3 | 0.1 | 0.2 | 1.4 | 5.8 | 0.5 | 1.9 | 6.3 | 7.8 | 75.1 |
| CXN | VZ | 14.1 | 91.9 | 106.0 | 791.0 | 103.0 | 3.8 | 33.9 | 54.7 | 37.7 | 0.7 | 0.8 | 1.7 | 11.0 | 1.9 | 3.5 | 12.8 | 14.8 | 47.1 |
| CXN | VZ | 16.2 | 148.2 | 164.4 | 751.1 | 84.5 | 4.1 | 14.0 | 46.7 | 25.0 | 0.4 | 0.4 | 1.1 | 6.3 | 1.1 | 2.2 | 7.3 | 14.7 | 52.6 |
| CXN | VZ | 11.9 | 35.6 | 47.5 | 831.0 | 121.5 | 4.1 | 12.4 | 51.3 | 19.7 | 0.8 | 0.7 | 1.2 | 6.3 | 1.8 | 3.0 | 8.1 | 22.2 | 40.2 |

Table A2. Number of reads and ASVs per plot. Sample gives information about sample type, either soil, or litter sample. Flood levels are TF = terra firme (not seasonally flooded); HV = high várzea (flooded 0-1 mo/yr); MV = mid várzea (flooded 2-4 mo/yr); and LV = low várzea (flooded 5-12 mo/yr). Lat and Long inform about latitude and longitude. ASV 16S is the number of unique amplicon sequence variants for Prokaryotes within the V3-V4 region (~460 bases) of the 16S rDNA gene. Reads 16S is the number of read amplicon sequence variants for Prokaryotes using the forward primer (5’-CCTACGGGN GGCWGCAG-3’) and reverse primer (5’-GACTACH VGGGTATCTAATCC-3’) from Klindworth et al. (2013). ASV 18S is the number of unique amplicon sequence variants for Eukaryotes within the V7 region (100–110 bases) of the 18S rDNA gene. Reads 18S is the number of read amplicon sequence variants for Eukaryotes using the forward and reverse primers (5’-TTTGTCTGSTTAATTSCG-3’) and (5’-TCACAGACCTGTTATTGC-3’) designed by Guardiola et al. (2015).

| **ID** | **Sample** | **Flood level** | **Lat** | **Long** | **ASV 16S** | **Reads 16S** | **ASV 18S** | **Reads 18S** |
| --- | --- | --- | --- | --- | --- | --- | --- | --- |
| HV1L | Litter | HV | -5.95423 | -67.8206 | 3059 | 28032 | 1004 | 18669 |
| HV5L | Litter | HV | -5.73941 | -67.8002 | 4805 | 34181 | 76 | 15106 |
| HV5S | Soil | HV | -5.73941 | -67.8002 | 30 | 95 | 78 | 16603 |
| HV7S | Soil | HV | -5.62975 | -67.6728 | 70 | 84 | 46 | 11525 |
| HV8L | Litter | HV | -5.62829 | -67.6805 | 3418 | 24377 | 875 | 24647 |
| HV8S | Soil | HV | -5.62829 | -67.6805 | 81 | 202 | 60 | 10593 |
| LV10L | Litter | LV | -5.55946 | -67.5495 | 4477 | 23487 | 265 | 19092 |
| LV10S | Soil | LV | -5.55946 | -67.5495 | 105 | 680 | 455 | 19712 |
| LV13L | Litter | LV | -5.41871 | -67.525 | 4238 | 17393 | 521 | 20296 |
| LV13S | Soil | LV | -5.41871 | -67.525 | 1609 | 24327 | 112 | 19043 |
| LV14L | Litter | LV | -5.40669 | -67.2777 | 4496 | 35647 | 911 | 23505 |
| LV14S | Soil | LV | -5.40669 | -67.2777 | 1772 | 16793 | 57 | 4432 |
| LV15L | Litter | LV | -5.40663 | -67.2776 | 5389 | 47070 | 579 | 24503 |
| LV15S | Soil | LV | -5.40663 | -67.2776 | 3161 | 43808 | 582 | 12634 |
| LV16L | Litter | LV | -5.3906 | -67.2567 | 4788 | 42670 | 884 | 18191 |
| LV16S | Soil | LV | -5.3906 | -67.2567 | 3410 | 33305 | 276 | 14095 |
| LV4L | Litter | LV | -5.76675 | -67.7586 | 4292 | 38921 | 854 | 27067 |
| MV11L | Litter | MV | -5.55234 | -67.5464 | 4683 | 24436 | NA | NA |
| MV18L | Litter | MV | -5.10586 | -67.1315 | 3964 | 24003 | 918 | 18203 |
| MV2S | Soil | MV | -5.78612 | -67.8049 | 110 | 347 | 791 | 16971 |
| MV3L | Litter | MV | -5.78131 | -67.8081 | 5447 | 49443 | 52 | 8799 |
| MV6L | Litter | MV | -5.67433 | -67.7302 | 97 | 874 | 1183 | 20140 |
| TF12L | Litter | TF | -5.45701 | -67.2609 | 4269 | 29573 | 258 | 14856 |
| TF12S | Soil | TF | -5.45701 | -67.2609 | 4261 | 51260 | 863 | 18215 |
| TF17L | Litter | TF | -5.39057 | -67.2115 | 2890 | 35750 | 872 | 22443 |
| TF17S | Soil | TF | -5.39057 | -67.2115 | 2238 | 34608 | 577 | 12622 |
| TF19L | Litter | TF | -5.04703 | -67.1356 | 1225 | 23870 | 537 | 17475 |
| TF19S | Soil | TF | -5.04703 | -67.1356 | 219 | 1364 | 60 | 9321 |
| TF9L | Litter | TF | -5.57857 | -67.505 | 2045 | 32479 | 894 | 16777 |
| TF9S | Soil | TF | -5.57857 | -67.505 | 2147 | 38748 | 175 | 17892 |

Table A3. Proportional contribution of each physicochemical soil variable for the principal components (PC) of our principal components analysis. We also show the proportion of variance explained for each PC in Total and the cumulative of variance explained. Coarse, Fine, and Total sand indicate the content of sand with grainsize 2.0-0.2 mm, 0.2-0.05 mm or 2.0-0.05 mm, respectively, in g/kg. Silt and Clay indicate clay and silt content in g/kg. pH is that of sample when dissolved in water. P = phosphorus, K = potassium, and Na = Natrium are given in mg/dm^3^. Ca = Calcium, Mg = magnesium, Al = aluminium, H+Al = exchangeable aluminium, SB = sum of exchangeable bases, t = effective cation exchange capacity, T = cation exchange capacity, V = Base Saturation Index, and m = aluminium saturation index are given in cmol_c_/dm^3^.

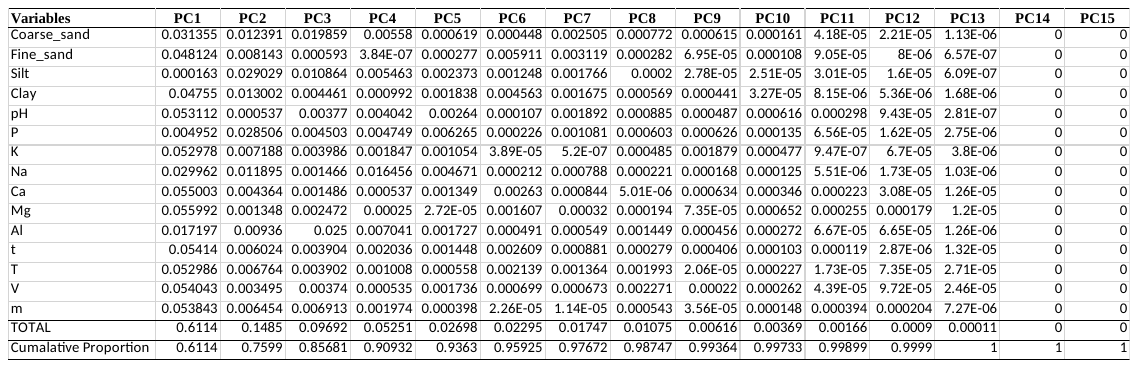

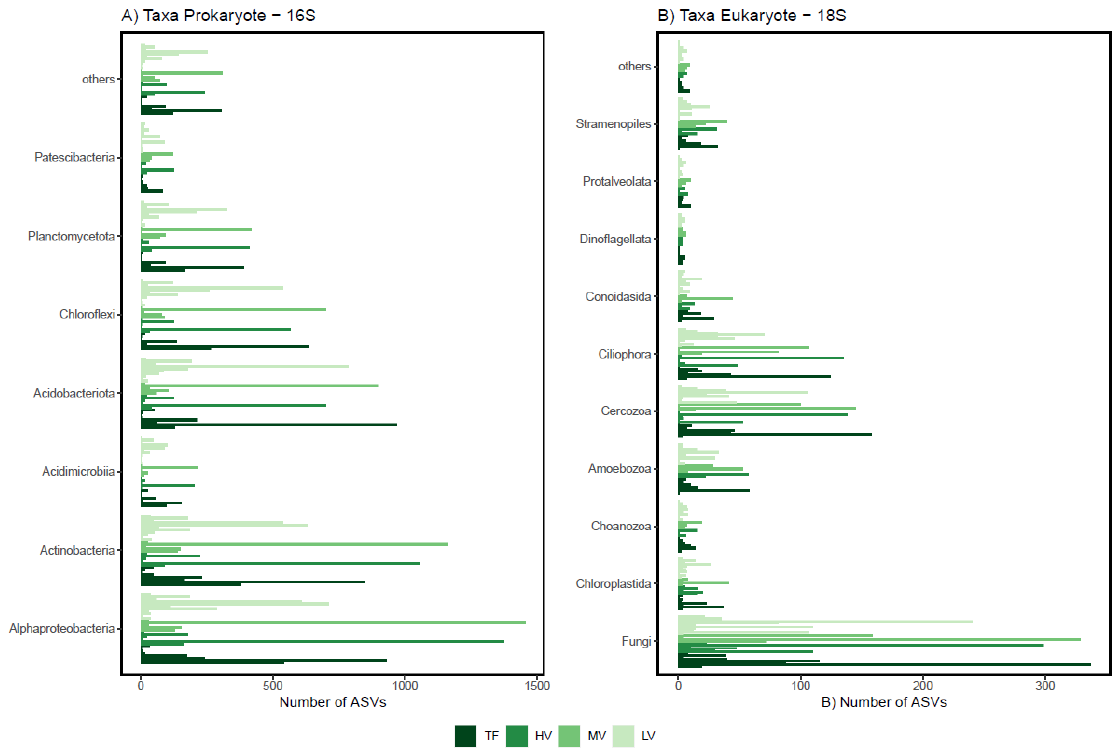

Fig. A1. **Taxonomic composition of amplicon sequence variants (ASV) communities.** Number of ASVs by taxonomic group and flood level for (A) prokaryotes and (B) eukaryotes. Flood levels are TF = terra firme (not seasonally flooded); HV = high várzea (flooded 0-1 mo/yr); MV = mid várzea (flooded 2-4 mo/yr); and LV = low várzea (flooded 5-12 mo/yr).

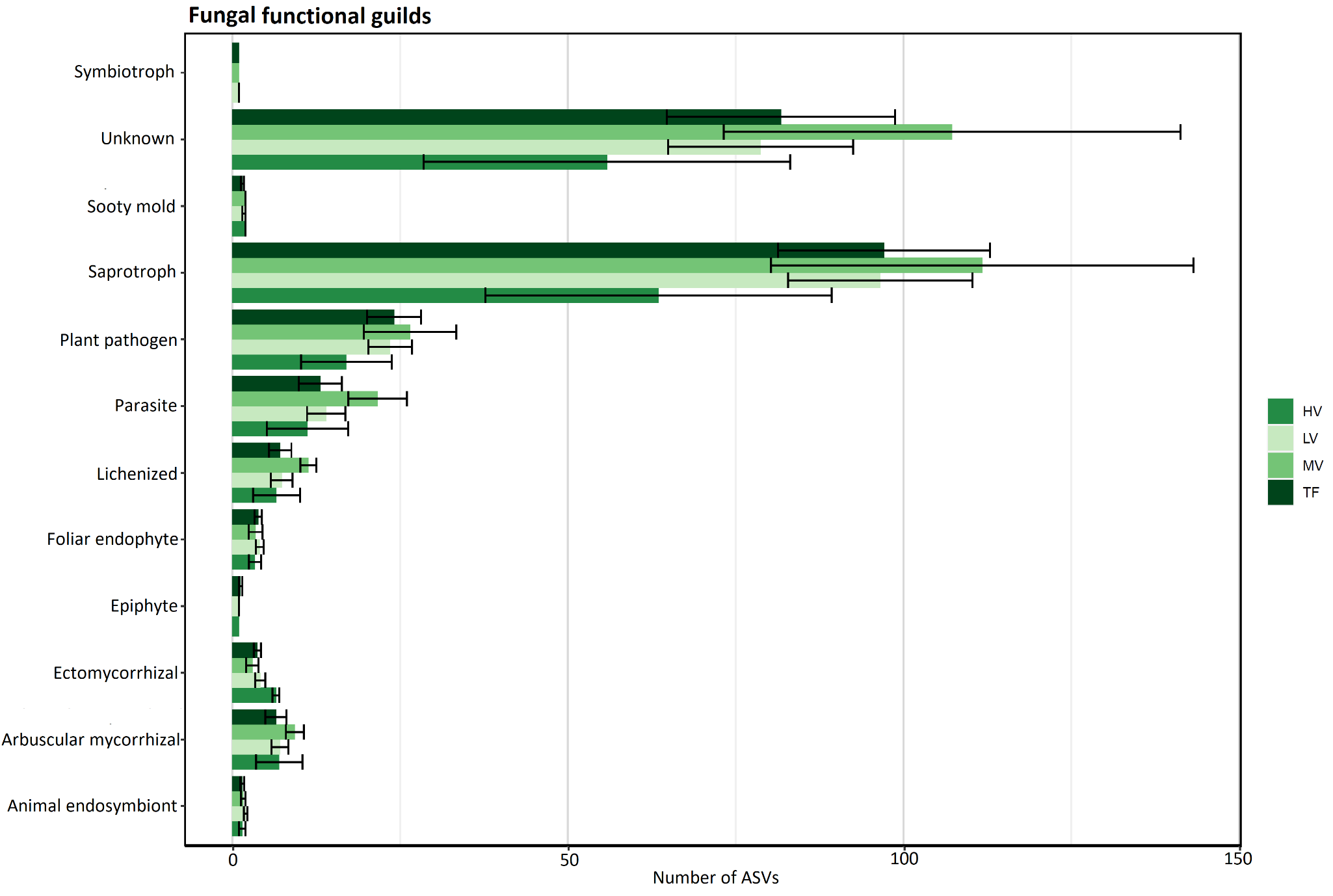

Fig. A2. Guild of fungi observed in the Juruá terra firme (TF; not seasonally flooded), high várzea (HV; flooded 0-1 mo/yr); mid várzea (MV; flooded 2-4 mo/yr); and low várzea (LV; flooded 5-12 mo/yr).
