## Appendix3 for "Above- and below-ground biodiversity responses to the prolonged flood pulse in central-western Amazonia, Brazil": paper2_Appendix3.html

Jurua soil metabarcoding Analysis


### Jurua soil metabarcoding Analysis

###### Camila Duarte Ritter

###### March 2021

```
##    ASV_ID TF12S TF17S LV15S LV14S LV16S MV2S HV5S HV8S HV7S TF19S TF9S LV10S
## 1 ASV9119     2     0     0     0     0    0    0    0    0     0    0     0
## 2 ASV9120     2     0     0     0     0    0    0    0    0     0    0     0
## 3 ASV9121     2     0     0     0     0    0    0    0    0     0    0     0
## 4 ASV9122     2     0     0     0     0    0    0    0    0     0    0     0
## 5 ASV9123     1     0     0     0     1    0    0    0    0     0    0     0
## 6 ASV9124     1     0     0     0     0    0    0    0    0     0    0     0
##   LV13S TF12L TF17L LV15L LV14L LV16L MV6L MV3L HV5L LV4L HV8L MV18L TF19L TF9L
## 1     0     0     0     0     0     0    0    0    0    0    0     0     0    0
## 2     0     0     0     0     0     0    0    0    0    0    0     0     0    0
## 3     0     0     0     0     0     0    0    0    0    0    0     0     0    0
## 4     0     0     0     0     0     0    0    0    0    0    0     0     0    0
## 5     0     0     0     0     0     0    0    0    0    0    0     0     0    0
## 6     0     0     0     0     0     1    0    0    0    0    0     0     0    0
##   MV11L LV10L LV13L HV1L
## 1     0     0     0    0
## 2     0     0     0    0
## 3     0     0     0    0
## 4     0     0     0    0
## 5     0     0     0    0
## 6     0     0     0    0
```

```
##                Community ID_Or    ID Sample forest  Code yen_tree_plot
## 1            Xibauazinho  LHV5  HV1L Litter     Vz  LVZ5  CAM-XZN-A-P5
## 2          Monte Carmelo  LHV8  HV5L Litter     Vz  LVZ8       McmB_P2
## 3             Morro Alto  LHV9  HV8L Litter     Vz  LVZ9       MltB_P4
## 4 Santo Antonio do Brito LLV11 LV10L Litter     Vz LVZ11       SanB_P4
## 5 São Raimundo (SrmC_P1)  LLV3 LV13L Litter     Vz  LVZ3       SrmC_P1
## 6       Bauana (BojE_P2)  LLV1 LV14L Litter     Vz  LVZ5       BojE_P2
##         PC1 fisher.alpha Flood_class meanAGB Cred_2.5 Cred_97.5 meanAGC
## 1  4.531045     81.57209        H_Vz   39.20    29.20      53.6   18.40
## 2  2.397633     30.28956        H_Vz   14.20    10.60      18.7    6.70
## 3 -1.793359     62.66386        H_Vz   40.60    32.00      50.9   19.30
## 4 -2.440236     56.61093        L_Vz   34.90    27.30      44.8   16.50
## 5 -2.540168     79.70264        L_Vz   25.87    18.67      36.1   12.13
## 6 -2.493159     14.66328        L_Vz   12.20     8.20      17.3    5.70
##   Cred_2.5.1 Cred_97.5.1 PC1.1   BA watermark_count watermark_average_cm
## 1      13.40        25.3 -1.51 4.39              62                   87
## 2       5.10         9.0 -1.10 1.56              55                    1
## 3      15.10        24.5 -1.41 4.32              71                   54
## 4      12.50        21.7 -2.18 3.76              54                  225
## 5       8.67        16.9 -1.28 3.18              NA                  168
## 6       3.70         8.1 -0.96 1.54              30                  185
##   watermark_max_cm watermark_min_cm watermark_StdDev_cm       Lat      Long
## 1              140               43                  27 -5.954230 -67.82061
## 2               26                0                   4 -5.739410 -67.80020
## 3              200                0                  49 -5.628290 -67.68045
## 4              275              140                  31 -5.559458 -67.54945
## 5              250               93                  37 -5.418710 -67.52496
## 6              260              124                  42 -5.406690 -67.27769
```

```
##    ASV_ID TF12S TF17S LV15S LV14S LV16S MV2S HV5S HV8S HV7S TF19S TF9S LV10S
## 1 ASV1094     5     0     0     0     0    1    0    0    0     0    0     1
## 2 ASV1250     1     0     0     0     0    0    0    0    0     0    0     0
## 3 ASV1599     1     0     0     0     0    0    0    0    0     0    0     0
## 4 ASV1936     0     0     2     0     0    2    0    0    0     0    0     0
## 5 ASV1943     0     0     0     0     0    0    0    0    0     0    0     0
## 6 ASV2137     0     0     0     0     0    0    0    0    0     0    0     0
##   LV13S TF12L TF17L LV15L LV14L LV16L MV6L MV3L HV5L LV4L HV8L MV18L TF19L TF9L
## 1     0     0     1     0     1     6    0    0    0    5    5     2     0    0
## 2     0     0     2     0     0     0    0    0    0    0    1     8     0    2
## 3     0     0     0     1     1     0    1    0    0    2    0     0     0    3
## 4     0     0     0     0     0     0    0    0    0    1    0     0     0    0
## 5     0     0     0     0     0     0    0    0    0    0    0     2     0    2
## 6     0     0     0     0     0     1    0    0    0    0    1     0     0    1
##   LV10L LV13L HV1L
## 1     0     0    0
## 2     0     0    6
## 3     0     0    2
## 4     0     0    1
## 5     0     0    2
## 6     0     0    0
```

```
##                Community ID_Or    ID Sample forest  Code yen_tree_plot
## 1            Xibauazinho  LHV5  HV1L Litter     Vz  LVZ5  CAM-XZN-A-P5
## 2          Monte Carmelo  LHV8  HV5L Litter     Vz  LVZ8       McmB_P2
## 3             Morro Alto  LHV9  HV8L Litter     Vz  LVZ9       MltB_P4
## 4 Santo Antonio do Brito LLV11 LV10L Litter     Vz LVZ11       SanB_P4
## 5 São Raimundo (SrmC_P1)  LLV3 LV13L Litter     Vz  LVZ3       SrmC_P1
## 6       Bauana (BojE_P2)  LLV1 LV14L Litter     Vz  LVZ5       BojE_P2
##         PC1 fisher.alpha Flood_class meanAGB Cred_2.5 Cred_97.5 meanAGC
## 1  4.531045     81.57209        H_Vz   39.20    29.20      53.6   18.40
## 2  2.397633     30.28956        H_Vz   14.20    10.60      18.7    6.70
## 3 -1.793359     62.66386        H_Vz   40.60    32.00      50.9   19.30
## 4 -2.440236     56.61093        L_Vz   34.90    27.30      44.8   16.50
## 5 -2.540168     79.70264        L_Vz   25.87    18.67      36.1   12.13
## 6 -2.493159     14.66328        L_Vz   12.20     8.20      17.3    5.70
##   Cred_2.5.1 Cred_97.5.1 PC1.1   BA watermark_count watermark_average_cm
## 1      13.40        25.3 -1.51 4.39              62                   87
## 2       5.10         9.0 -1.10 1.56              55                    1
## 3      15.10        24.5 -1.41 4.32              71                   54
## 4      12.50        21.7 -2.18 3.76              54                  225
## 5       8.67        16.9 -1.28 3.18              NA                  168
## 6       3.70         8.1 -0.96 1.54              30                  185
##   watermark_max_cm watermark_min_cm watermark_StdDev_cm       Lat      Long
## 1              140               43                  27 -5.954230 -67.82061
## 2               26                0                   4 -5.739410 -67.80020
## 3              200                0                  49 -5.628290 -67.68045
## 4              275              140                  31 -5.559458 -67.54945
## 5              250               93                  37 -5.418710 -67.52496
## 6              260              124                  42 -5.406690 -67.27769
```

```
####### ###############

soil_phy_che <- read.csv("input/Soil_metadata.csv", header = T)
head(soil_phy_che)
```

```
##   Locality habitat ID_org  Plot Plot_org Flood_class Forest Coarse_sand
## 1    Jurua    J_TF   LHV5  HV1W  BauA_P3        J_TF     TF      13.670
## 2    Jurua    J_TF   LHV8  HV5W  BojA_P1        J_TF     TF      39.380
## 3    Jurua    L_Vz   LHV9  HV8W  BojE_P2        L_Vz     Vz       5.940
## 4    Jurua    L_Vz   LLV1 LV14W  BojE_P5        L_Vz     Vz      11.255
## 5    Jurua    M_Vz  LLV11 LV10W  CarD_P1        M_Vz     Vz      17.375
## 6    Jurua    M_Vz  LLV15  LV4W  ItaB_P2        M_Vz     Vz       4.030
##   Fine_sand total_sand_fraction    Silt  Clay   pH         P   K Na    Ca   Mg
## 1   418.480             432.160 426.340 141.5 4.04  5.000000  46  5  0.09 0.12
## 2   129.320             168.700 382.800 448.5 4.00  3.000000  86 45  0.38 0.44
## 3    12.910              18.850 512.650 468.5 4.67 10.808030 189 28  9.12 2.84
## 4    15.025              26.280 275.720 698.0 4.65  7.398785 146 42 10.59 3.62
## 5    15.920              33.295 314.205 652.5 4.63  6.310729 142 34 11.76 3.70
## 6     9.295              13.325 441.175 545.5 4.61  8.486842 145 35 13.89 3.11
##     Al   H_Al       SB        t        T        V        m
## 1 3.59  7.760  0.35000  3.94000  8.10000  4.31000 91.13000
## 2 8.17 11.700  1.24000  9.41000 12.93000  9.55000 86.86000
## 3 2.09  8.019 12.56512 14.65512 20.58412 61.04278 14.26123
## 4 1.85 10.758 14.76601 16.61601 25.52401 57.85145 11.13384
## 5 1.93  9.768 15.97100 17.90100 25.73900 62.04980 10.78152
## 6 3.83  9.339 17.52302 21.35302 26.86202 65.23344 17.93657
```

```
# Standarzi the variables
ncol(soil_phy_che)
```

```
## [1] 25
```

```
soil_phy_che_st <- scale(soil_phy_che[, 8:25])
head(soil_phy_che_st)
```

```
##      Coarse_sand  Fine_sand total_sand_fraction        Silt        Clay
## [1,]  -0.3153884  2.7545422          1.15981519  0.07230477 -1.30472817
## [2,]  -0.1519772  0.3338590          0.06628333 -0.14366914  0.05766632
## [3,]  -0.3645198 -0.6406594         -0.55569250  0.50043312  0.14642166
## [4,]  -0.3307379 -0.6229538         -0.52485312 -0.67482408  1.16488920
## [5,]  -0.2918396 -0.6154614         -0.49573627 -0.48392477  0.96297080
## [6,]  -0.3766596 -0.6709221         -0.57862487  0.14589165  0.48812972
##              pH           P          K         Na         Ca         Mg
## [1,] -0.7977447 -0.77062189 -1.1703000 -1.8605381 -1.1520174 -1.3135492
## [2,] -0.8936179 -1.02244754 -0.4105020  1.7424405 -1.1068789 -1.1096897
## [3,]  0.7122578 -0.03931649  1.5459779  0.2111746  0.2535019  0.4192570
## [4,]  0.6643212 -0.46858405  0.7291950  1.4722171  0.4823074  0.9161646
## [5,]  0.6163847 -0.60558434  0.6532152  0.7516214  0.6644179  0.9671295
## [6,]  0.5684481 -0.33158377  0.7102001  0.8416958  0.9959524  0.5912635
##              Al        H_Al         SB          t          T          V
## [1,]  0.6614366 -0.13380548 -1.2017515 -1.1039112 -1.1298930 -1.4159155
## [2,]  3.3084671  1.22839317 -1.0916005 -0.3989431 -0.5898938 -1.2319991
## [3,] -0.2054949 -0.04425993  0.3100547  0.2770420  0.2658445  0.5753227
## [4,] -0.3442039  0.90271014  0.5824489  0.5297602  0.8181300  0.4633119
## [5,] -0.2979675  0.56043180  0.7315844  0.6953681  0.8421658  0.6106680
## [6,]  0.8001456  0.41211119  0.9236705  1.1402610  0.9677207  0.7224090
##               m
## [1,]  1.7333832
## [2,]  1.6008551
## [3,] -0.6523938
## [4,] -0.7494587
## [5,] -0.7603936
## [6,] -0.5383222
```

```
soil_phy_che_st <- cbind(soil_phy_che[1:7], soil_phy_che_st)

# Principle component analyses

## All
All_soil <- c("Coarse_sand", "Fine_sand", "Silt", "Clay", "pH", "P", "K", "Na", "Ca", 
    "Mg", "Al", "t", "T", "V", "m")

All_soil.pca <- prcomp(soil_phy_che_st[, All_soil], center = F, scale. = F)

# Making the percetagem
load <- with(All_soil.pca, unclass(rotation))
load
```

```
##                       PC1         PC2         PC3           PC4          PC5
## Coarse_sand  0.1806560624 -0.24871402  0.54852453 -2.677269e-01 -0.065614832
## Fine_sand    0.2772693449 -0.16344990  0.01638484 -1.843856e-05  0.029397195
## Silt         0.0009413444  0.58268037 -0.30007107  2.621368e-01  0.251530904
## Clay        -0.2739606772 -0.26099135 -0.12321015 -4.758243e-02 -0.194799662
## pH          -0.3060057797  0.01078618  0.10411826  1.939244e-01  0.279837864
## P           -0.0285330085  0.57218647  0.12438666 -2.278641e-01 -0.664100416
## K           -0.3052376776 -0.14429013 -0.11009098  8.863920e-02  0.111771643
## Na          -0.1726274296  0.23875251  0.04049039 -7.896221e-01  0.495117324
## Ca          -0.3169065646 -0.08759452  0.04105177 -2.577846e-02 -0.142945876
## Mg          -0.3226024134 -0.02705651  0.06827904  1.201623e-02  0.002888331
## Al           0.0990796101 -0.18787348 -0.69052820 -3.378383e-01 -0.183079787
## t           -0.3119338869 -0.12090660 -0.10784504 -9.767781e-02 -0.153473503
## T           -0.3052801669 -0.13576478 -0.10777744 -4.836861e-02 -0.059115507
## V           -0.3113703677  0.07016052  0.10330563  2.569133e-02 -0.184005852
## m            0.3102204486 -0.12954608 -0.19094973 -9.470824e-02 -0.042144411
##                      PC6           PC7          PC8          PC9        PC10
## Coarse_sand -0.050343306 -4.462800e-01 -0.205615873  0.259796590 -0.13345380
## Fine_sand   -0.664073228  5.557230e-01  0.075117454  0.029354459  0.08947753
## Silt        -0.140162010 -3.146198e-01 -0.053246180 -0.011744944  0.02081290
## Clay         0.512587252  2.984771e-01  0.151373658 -0.186434651  0.02712041
## pH          -0.011964617  3.371126e-01 -0.235714421  0.205704253 -0.51096586
## P           -0.025390610  1.925836e-01  0.160423707  0.264779003 -0.11183373
## K            0.004364705  9.257186e-05  0.128995095  0.794298469  0.39611770
## Na           0.023768149  1.404025e-01  0.058776362 -0.071122411  0.10356162
## Ca          -0.295428030 -1.503684e-01 -0.001333821 -0.267925791  0.28754230
## Mg          -0.180507050 -5.700637e-02  0.051648791 -0.031062887 -0.54110926
## Al          -0.055167675 -9.775649e-02 -0.385785015  0.192782482 -0.22583538
## t           -0.293106344 -1.568821e-01 -0.074191068 -0.171746246  0.08561955
## T           -0.240244515 -2.430489e-01  0.530536704 -0.008709289 -0.18840000
## V           -0.078550811  1.199167e-01 -0.604640818 -0.092971830  0.21756773
## m           -0.002536660 -2.029083e-03  0.144483252 -0.015066681 -0.12306753
##                     PC11         PC12         PC13          PC14          PC15
## Coarse_sand  0.066471396  0.067074280  0.026029430  6.455192e-03 -4.355727e-01
## Fine_sand   -0.143982871  0.024287336  0.015067613  4.827213e-03 -3.307114e-01
## Silt         0.047908536 -0.048560252  0.013979273  8.243777e-03 -5.581241e-01
## Clay        -0.012957413 -0.016272297 -0.038627280  9.265445e-03 -6.238470e-01
## pH           0.474129001  0.286178220  0.006448157  1.994341e-04  2.147163e-06
## P            0.104388783  0.049151457 -0.062996878  2.930554e-05 -8.443866e-07
## K           -0.001506079 -0.203158810 -0.087272860  1.303999e-02  1.955370e-04
## Na           0.008757442 -0.052388367  0.023535769  4.718997e-03  7.165351e-05
## Ca           0.355501654  0.093516495 -0.289090057  6.212690e-01  9.174671e-03
## Mg          -0.405744444 -0.543102598 -0.274694760  1.518729e-01  2.246177e-03
## Al          -0.106075019  0.201780195 -0.028855765  1.673384e-01  2.475907e-03
## t            0.188878775  0.008708129 -0.303443171 -7.500306e-01 -1.110771e-02
## T           -0.027511925  0.223174268  0.621713899 -3.921603e-04  5.632462e-06
## V           -0.069833409 -0.294953253  0.564748868 -3.049172e-04  1.219776e-05
## m            0.626930924 -0.618591446  0.166784706  2.295466e-05  3.139778e-06
```

```
PCprop <- abs(load)  ## save absolute values
PCprop2 <- sweep(PCprop, 2, colSums(PCprop), "/")
PCprop2 <- PCprop2 * 100
write.csv(PCprop2, "PCprop.csv")

write.csv(All_soil.pca$rotation, "All_soil.pca.csv")

screeplot(All_soil.pca)
```

```
summary(All_soil.pca)
```

```
## Importance of components:
##                           PC1    PC2     PC3     PC4     PC5     PC6     PC7
## Standard deviation     3.0284 1.4923 1.20576 0.88753 0.63616 0.58670 0.51192
## Proportion of Variance 0.6114 0.1485 0.09692 0.05251 0.02698 0.02295 0.01747
## Cumulative Proportion  0.6114 0.7599 0.85681 0.90932 0.93630 0.95925 0.97672
##                            PC8     PC9    PC10    PC11   PC12    PC13      PC14
## Standard deviation     0.40164 0.30406 0.23530 0.15787 0.1159 0.04140 5.463e-05
## Proportion of Variance 0.01075 0.00616 0.00369 0.00166 0.0009 0.00011 0.000e+00
## Cumulative Proportion  0.98747 0.99364 0.99733 0.99899 0.9999 1.00000 1.000e+00
##                             PC15
## Standard deviation     4.145e-06
## Proportion of Variance 0.000e+00
## Cumulative Proportion  1.000e+00
```

```
write.csv(All_soil.pca$x, "All_soil.pca2.csv")
```

```
cols <- c("#238b45", "#00441b", "#99d8c9", "#41ae76", "#762a83", "#c2a5cf")


FigPCA_All <- autoplot(All_soil.pca, data = soil_phy_che_st, colour = "Flood_class", 
    shape = "Locality", size = 3, asp = 1, loadings = T, frame = T, loadings.colour = "gray", 
    plot.background = element_blank(), loadings.label = T, loadings.label.size = 5, 
    loadings.label.colour = "black") + theme(plot.background = element_blank(), panel.background = element_rect(fill = "transparent", 
    color = "black", size = 1), legend.text = element_text(), legend.key = element_blank()) + 
    scale_fill_manual(values = cols) + scale_color_manual(values = cols) + scale_size_continuous(range = c(5, 
    15))
```

```
## Warning: `select_()` is deprecated as of dplyr 0.7.0.
## Please use `select()` instead.
## This warning is displayed once every 8 hours.
## Call `lifecycle::last_warnings()` to see where this warning was generated.
```

```
## Warning: `group_by_()` is deprecated as of dplyr 0.7.0.
## Please use `group_by()` instead.
## See vignette('programming') for more help
## This warning is displayed once every 8 hours.
## Call `lifecycle::last_warnings()` to see where this warning was generated.
```

```
ggsave("FigPCA_All.pdf", plot = FigPCA_All, width = 12, height = 8)
FigPCA_All
```

```
##### Diversity Estimates########

# 16S

plots_16S <- read.csv("input/metadata_16S.csv")

rownames(plots_16S) <- plots_16S$ID
plots_16S <- plots_16S %>% select(-ID)

abundance_16S <- phyloseq(otu_table(ASVs_M16S, taxa_are_rows = TRUE), sample_data(plots_16S))

# 18S

plots_18S <- read.csv("input/metadata_18S.csv")

rownames(plots_18S) <- plots_18S$ID
plots_18S <- plots_18S %>% select(-ID)

abundance_18S <- phyloseq(otu_table(ASVs_M18S, taxa_are_rows = TRUE), sample_data(plots_18S))

# Figure

colsSample <- c("#4d004b", "#8c96c6")

colsFlood <- c("#00441b", "#238b45", "#74c476", "#c7e9c0")

Fig_Div_16S <- plot_richness(abundance_16S, x = "Flood_class", shape = "Sample", 
    measures = c("Observed", "Chao1", "Fisher"), color = "Flood_class") + scale_color_manual(values = colsFlood) + 
    geom_point(size = 3) + ggtitle("A) Alpha diversity - 16S") + ylab("") + xlab("") + 
    theme(panel.grid.major.x = element_line(size = 0.5, colour = "#f0f0f0"), panel.grid.major.y = element_blank(), 
        panel.grid.minor = element_blank(), panel.background = element_blank(), axis.line = element_line(colour = "black"), 
        panel.border = element_rect(colour = "black", fill = NA, size = 1), legend.position = ("none"))


Fig_Div_18S <- plot_richness(abundance_18S, x = "Flood_class", shape = "Sample", 
    measures = c("Observed", "Chao1", "Fisher"), color = "Flood_class") + scale_color_manual(values = colsFlood) + 
    geom_point(size = 3) + ggtitle("B) Alpha diversity - 18S") + ylab("") + xlab("") + 
    theme(panel.grid.major.x = element_line(size = 0.5, colour = "#f0f0f0"), panel.grid.major.y = element_blank(), 
        panel.grid.minor = element_blank(), panel.background = element_blank(), axis.line = element_line(colour = "black"), 
        panel.border = element_rect(colour = "black", fill = NA, size = 1), legend.position = ("none"))

newSOrder = c("TF", "H_Vz", "M_Vz", "L_Vz")

Fig_Div_16S$data$Flood_class <- as.character(Fig_Div_16S$data$Flood_class)
Fig_Div_16S$data$Flood_class <- factor(Fig_Div_16S$data$Flood_class, levels = newSOrder)

Fig_Div_18S$data$Flood_class <- as.character(Fig_Div_18S$data$Flood_class)
Fig_Div_18S$data$Flood_class <- factor(Fig_Div_18S$data$Flood_class, levels = newSOrder)

figAlphaDiv <- ggarrange(Fig_Div_16S, Fig_Div_18S, common.legend = TRUE, legend = "bottom", 
    ncol = 2)
ggsave("fig_Alpha_Diversity.pdf", plot = figAlphaDiv, width = 9, height = 6)
figAlphaDiv
```

```
#### Create Table with diversity and metadata_16S

Alpha_data_16S <- estimate_richness(abundance_16S, split = TRUE, measures = c("Observed", 
    "Chao1", "Shannon", "Simpson", "Fisher"))

Alpha_data_16S <- Alpha_data_16S %>% rownames_to_column(var = "ID")

Alpha_data_16S <- full_join(metadata_16S, Alpha_data_16S, by = "ID")
Alpha_data_16S_st <- scale(Alpha_data_16S[, c("Fisher", "watermark_max_cm", "fisher.alpha")])
Alpha_data_16S_st2 <- cbind(Alpha_data_16S[, c("ID", "Flood_class", "Sample", "PC1")], 
    Alpha_data_16S_st)

### Model selection for explanatory variables

m0_16S = glm(Fisher ~ 1, data = Alpha_data_16S_st2)
m1_16S = glm(Fisher ~ Flood_class, data = Alpha_data_16S_st2)
m2_16S = glm(Fisher ~ Sample, data = Alpha_data_16S_st2)
m3_16S = glm(Fisher ~ watermark_max_cm, data = Alpha_data_16S_st2)
m4_16S = glm(Fisher ~ PC1, data = Alpha_data_16S_st2)
m5_16S = glm(Fisher ~ PC1 * Flood_class, data = Alpha_data_16S_st2)
m6_16S = glm(Fisher ~ fisher.alpha, data = Alpha_data_16S_st2)
m7_16S = glm(Fisher ~ fisher.alpha * Flood_class, data = Alpha_data_16S_st2)
m8_16S = glm(Fisher ~ fisher.alpha * Sample, data = Alpha_data_16S_st2)


AICctab(m0_16S, m1_16S, m2_16S, m3_16S, m4_16S, m5_16S, m6_16S, m7_16S, m8_16S, base = TRUE, 
    nobs = 16, weights = TRUE)
```

```
##        AICc  dAICc df weight
## m8_16S  73.1   0.0 5  0.7625
## m2_16S  75.5   2.3 3  0.2355
## m6_16S  85.6  12.5 3  0.0015
## m0_16S  89.0  15.9 2  <0.001
## m3_16S  89.7  16.5 3  <0.001
## m4_16S  91.9  18.7 3  <0.001
## m1_16S  96.5  23.4 5  <0.001
## m7_16S 113.3  40.1 9  <0.001
## m5_16S 127.2  54.0 9  <0.001
```

```
summary(m8_16S)
```

```
## 
## Call:
## glm(formula = Fisher ~ fisher.alpha * Sample, data = Alpha_data_16S_st2)
## 
## Deviance Residuals: 
##      Min        1Q    Median        3Q       Max  
## -2.04241  -0.39905   0.03446   0.49400   1.00575  
## 
## Coefficients:
##                         Estimate Std. Error t value Pr(>|t|)    
## (Intercept)               0.6427     0.1815   3.540  0.00167 ** 
## fisher.alpha             -0.2288     0.1947  -1.175  0.25150    
## SampleSoil               -1.3463     0.2774  -4.853 6.03e-05 ***
## fisher.alpha:SampleSoil   0.4439     0.2797   1.587  0.12566    
## ---
## Signif. codes:  0 '***' 0.001 '**' 0.01 '*' 0.05 '.' 0.1 ' ' 1
## 
## (Dispersion parameter for gaussian family taken to be 0.5256352)
## 
##     Null deviance: 26.317  on 27  degrees of freedom
## Residual deviance: 12.615  on 24  degrees of freedom
##   (2 observations deleted due to missingness)
## AIC: 67.136
## 
## Number of Fisher Scoring iterations: 2
```

```
#### Create Table with diversity and metadata_18S

Alpha_data_18S <- estimate_richness(abundance_18S, split = TRUE, measures = c("Observed", 
    "Chao1", "Shannon", "Simpson", "Fisher"))

Alpha_data_18S <- Alpha_data_18S %>% rownames_to_column(var = "ID")

Alpha_data_18S <- full_join(metadata_18S, Alpha_data_18S, by = "ID")
Alpha_data_18S_st <- scale(Alpha_data_18S[, c("Fisher", "watermark_max_cm", "fisher.alpha")])
Alpha_data_18S_st2 <- cbind(Alpha_data_18S[, c("ID", "Flood_class", "Sample", "PC1")], 
    Alpha_data_18S_st)

### Model selection for explanatory variables

m0_18S = glm(Fisher ~ 1, data = Alpha_data_18S_st2)
m1_18S = glm(Fisher ~ Flood_class, data = Alpha_data_18S_st2)
m2_18S = glm(Fisher ~ Sample, data = Alpha_data_18S_st2)
m3_18S = glm(Fisher ~ watermark_max_cm, data = Alpha_data_18S_st2)
m4_18S = glm(Fisher ~ PC1, data = Alpha_data_16S_st2)
m5_18S = glm(Fisher ~ PC1 * Flood_class, data = Alpha_data_16S_st2)
m6_18S = glm(Fisher ~ fisher.alpha, data = Alpha_data_18S_st2)
m7_18S = glm(Fisher ~ fisher.alpha * Flood_class, data = Alpha_data_18S_st2)
m8_18S = glm(Fisher ~ fisher.alpha * Sample, data = Alpha_data_18S_st2)


AICctab(m0_18S, m1_18S, m2_18S, m3_18S, m4_18S, m5_18S, m6_18S, m7_18S, m8_18S, base = TRUE, 
    nobs = 16, weights = TRUE)
```

```
##        AICc  dAICc df weight
## m2_18S  82.4   0.0 3  0.4357
## m6_18S  83.6   1.1 3  0.2454
## m8_18S  83.7   1.3 5  0.2328
## m0_18S  86.2   3.8 2  0.0660
## m3_18S  89.2   6.8 3  0.0148
## m4_18S  91.9   9.4 3  0.0039
## m1_18S  94.0  11.6 5  0.0013
## m7_18S 111.7  29.3 9  <0.001
## m5_18S 127.2  44.7 9  <0.001
```

```
summary(m6_18S)
```

```
## 
## Call:
## glm(formula = Fisher ~ fisher.alpha, data = Alpha_data_18S_st2)
## 
## Deviance Residuals: 
##     Min       1Q   Median       3Q      Max  
## -1.2308  -1.0536  -0.1407   0.8197   2.0187  
## 
## Coefficients:
##              Estimate Std. Error t value Pr(>|t|)
## (Intercept)   0.04326    0.19617   0.221    0.827
## fisher.alpha  0.15527    0.19990   0.777    0.445
## 
## (Dispersion parameter for gaussian family taken to be 1.039004)
## 
##     Null deviance: 26.602  on 26  degrees of freedom
## Residual deviance: 25.975  on 25  degrees of freedom
##   (2 observations deleted due to missingness)
## AIC: 81.578
## 
## Number of Fisher Scoring iterations: 2
```

```
colsFlood <- c("#238b45", "#c7e9c0", "#74c476", "#00441b")

FigFisher_16S <- ggplot(Alpha_data_16S, aes(x = ID)) + geom_bar(aes(y = fisher.alpha, 
    colour = Flood_class, fill = Flood_class), stat = "identity", position = "dodge") + 
    geom_bar(aes(y = -1 * Fisher, colour = Flood_class, fill = Flood_class), stat = "identity", 
        position = "dodge") + scale_color_manual(values = colsFlood) + scale_fill_manual(values = colsFlood) + 
    geom_hline(yintercept = 0, size = 1) + ggtitle("A) Fisher diversity - 16S") + 
    ylab("Fisher") + xlab("") + ylim(-1800, 300) + theme(legend.position = "none") + 
    theme(axis.text.x = element_blank(), axis.ticks.x = element_blank(), panel.grid.major = element_blank(), 
        panel.grid.minor = element_blank(), panel.background = element_blank(), axis.line = element_line(colour = "black"), 
        panel.border = element_rect(colour = "black", fill = NA, size = 1))

FigFisher_18S <- ggplot(Alpha_data_18S, aes(x = ID)) + geom_bar(aes(y = fisher.alpha, 
    colour = Flood_class, fill = Flood_class), stat = "identity", position = "dodge") + 
    geom_bar(aes(y = -1 * Fisher, colour = Flood_class, fill = Flood_class), stat = "identity", 
        position = "dodge") + scale_color_manual(values = colsFlood) + scale_fill_manual(values = colsFlood) + 
    geom_hline(yintercept = 0, size = 1) + ggtitle("B) Fisher diversity - 18S") + 
    ylab("Fisher") + xlab("") + ylim(-1800, 300) + theme(legend.position = "none") + 
    theme(axis.text.x = element_blank(), axis.ticks.x = element_blank(), panel.grid.major = element_blank(), 
        panel.grid.minor = element_blank(), panel.background = element_blank(), axis.line = element_line(colour = "black"), 
        panel.border = element_rect(colour = "black", fill = NA, size = 1))

newSOrder2 = c("TF12L", "TF12S", "TF17L", "TF17S", "TF9S", "TF9L", "TF19L", "TF19S", 
    "HV1L", "HV7S", "HV8L", "HV8S", "HV5L", "HV5S", "MV2S", "MV6L", "MV18L", "MV11L", 
    "MV3L", "LV13L", "LV13S", "LV10L", "LV10S", "LV16L", "LV16S", "LV4L", "LV15L", 
    "LV15S", "LV14L", "LV14S")

FigFisher_16S$data$ID <- as.character(FigFisher_16S$data$ID)
FigFisher_16S$data$ID <- factor(FigFisher_16S$data$ID, levels = newSOrder2)

FigFisher_18S$data$ID <- as.character(FigFisher_18S$data$ID)
FigFisher_18S$data$ID <- factor(FigFisher_18S$data$ID, levels = newSOrder2)

figFisher <- ggarrange(FigFisher_16S, FigFisher_18S, common.legend = TRUE, legend = "bottom", 
    ncol = 2)
```

```
## Warning: Removed 2 rows containing missing values (geom_bar).

## Warning: Removed 2 rows containing missing values (geom_bar).

## Warning: Removed 2 rows containing missing values (geom_bar).
```

```
ggsave("FigBelowXAboveFisher.pdf", plot = figFisher, width = 9, height = 6)
figFisher
```

```
## envfit

### Eukaryote(16S)
envfit(sc16S2[, 1:2] ~ as.factor(sc16S2$forest))
```

```
## 
## ***FACTORS:
## 
## Centroids:
##                              NMDS1   NMDS2
## as.factor(sc16S2$forest)Tf -0.6891 -0.8078
## as.factor(sc16S2$forest)Vz  0.2506  0.2937
## 
## Goodness of fit:
##                              r2 Pr(>r)  
## as.factor(sc16S2$forest) 0.1237  0.026 *
## ---
## Signif. codes:  0 '***' 0.001 '**' 0.01 '*' 0.05 '.' 0.1 ' ' 1
## Permutation: free
## Number of permutations: 999
```

```
envfit(sc16S2[, 1:2] ~ as.factor(sc16S2$Flood_class))
```

```
## 
## ***FACTORS:
## 
## Centroids:
##                                     NMDS1   NMDS2
## as.factor(sc16S2$Flood_class)H_Vz  1.1350  0.8362
## as.factor(sc16S2$Flood_class)L_Vz -0.3418  0.0000
## as.factor(sc16S2$Flood_class)M_Vz  0.4926  0.2891
## as.factor(sc16S2$Flood_class)TF   -0.6891 -0.8078
## 
## Goodness of fit:
##                                 r2 Pr(>r)  
## as.factor(sc16S2$Flood_class) 0.24  0.025 *
## ---
## Signif. codes:  0 '***' 0.001 '**' 0.01 '*' 0.05 '.' 0.1 ' ' 1
## Permutation: free
## Number of permutations: 999
```

```
envfit(sc16S2[, 1:2] ~ as.factor(sc16S2$Sample))
```

```
## 
## ***FACTORS:
## 
## Centroids:
##                                  NMDS1   NMDS2
## as.factor(sc16S2$Sample)Litter -0.7101  0.3765
## as.factor(sc16S2$Sample)Soil    0.9285 -0.4924
## 
## Goodness of fit:
##                              r2 Pr(>r)    
## as.factor(sc16S2$Sample) 0.2549  0.001 ***
## ---
## Signif. codes:  0 '***' 0.001 '**' 0.01 '*' 0.05 '.' 0.1 ' ' 1
## Permutation: free
## Number of permutations: 999
```

```
## envfit

### Eukaryote(18S)
envfit(sc18S2[, 1:2] ~ as.factor(sc18S2$forest))
```

```
## 
## ***FACTORS:
## 
## Centroids:
##                              NMDS1   NMDS2
## as.factor(sc18S2$forest)Tf -0.1808 -0.2141
## as.factor(sc18S2$forest)Vz  0.0689  0.0816
## 
## Goodness of fit:
##                             r2 Pr(>r)
## as.factor(sc18S2$forest) 0.025   0.45
## Permutation: free
## Number of permutations: 999
```

```
envfit(sc18S2[, 1:2] ~ as.factor(sc18S2$Flood_class))
```

```
## 
## ***FACTORS:
## 
## Centroids:
##                                     NMDS1   NMDS2
## as.factor(sc18S2$Flood_class)H_Vz  0.7286  0.1558
## as.factor(sc18S2$Flood_class)L_Vz -0.1405  0.0408
## as.factor(sc18S2$Flood_class)M_Vz -0.3450  0.0823
## as.factor(sc18S2$Flood_class)TF   -0.1808 -0.2141
## 
## Goodness of fit:
##                                   r2 Pr(>r)
## as.factor(sc18S2$Flood_class) 0.1354  0.264
## Permutation: free
## Number of permutations: 999
```

```
envfit(sc18S2[, 1:2] ~ as.factor(sc18S2$Sample))
```

```
## 
## ***FACTORS:
## 
## Centroids:
##                                  NMDS1   NMDS2
## as.factor(sc18S2$Sample)Litter -0.4374  0.1456
## as.factor(sc18S2$Sample)Soil    0.5383 -0.1792
## 
## Goodness of fit:
##                              r2 Pr(>r)   
## as.factor(sc18S2$Sample) 0.2186  0.008 **
## ---
## Signif. codes:  0 '***' 0.001 '**' 0.01 '*' 0.05 '.' 0.1 ' ' 1
## Permutation: free
## Number of permutations: 999
```

```
## envfit

### Plants
envfit(plants2[, 1:2] ~ as.factor(plants2$forest))
```

```
## 
## ***FACTORS:
## 
## Centroids:
##                               NMDS1   NMDS2
## as.factor(plants2$forest)TF -1.0929 -0.0522
## as.factor(plants2$forest)Vz  0.2342  0.0112
## 
## Goodness of fit:
##                               r2 Pr(>r)   
## as.factor(plants2$forest) 0.4859  0.003 **
## ---
## Signif. codes:  0 '***' 0.001 '**' 0.01 '*' 0.05 '.' 0.1 ' ' 1
## Permutation: free
## Number of permutations: 999
```

```
envfit(plants2[, 1:2] ~ as.factor(plants2$Flood_class))
```

```
## 
## ***FACTORS:
## 
## Centroids:
##                                      NMDS1   NMDS2
## as.factor(plants2$Flood_class)H_Vz  0.1142  0.2606
## as.factor(plants2$Flood_class)L_Vz  0.1550 -0.3592
## as.factor(plants2$Flood_class)M_Vz  0.4175  0.0581
## as.factor(plants2$Flood_class)TF   -1.0929 -0.0522
## 
## Goodness of fit:
##                                    r2 Pr(>r)    
## as.factor(plants2$Flood_class) 0.6124  0.001 ***
## ---
## Signif. codes:  0 '***' 0.001 '**' 0.01 '*' 0.05 '.' 0.1 ' ' 1
## Permutation: free
## Number of permutations: 999
```

```
# 16S

tcom16S <- t(Abu_VST_16S3)
tcom16S <- as.data.frame(tcom16S)


ado16S <- adonis2(tcom16S ~ biol_16S$Flood_class * biol_16S$Sample, data = tcom16S, 
    type = "bray", binary = FALSE, permutations = 999, by = NULL)
ado16S
```

```
## Permutation test for adonis under reduced model
## Permutation: free
## Number of permutations: 999
## 
## adonis2(formula = tcom16S ~ biol_16S$Flood_class * biol_16S$Sample, data = tcom16S, permutations = 999, by = NULL, type = "bray", binary = FALSE)
##          Df SumOfSqs     R2      F Pr(>F)   
## Model     7   3.4899 0.3508 1.6983  0.003 **
## Residual 22   6.4584 0.6492                 
## Total    29   9.9483 1.0000                 
## ---
## Signif. codes:  0 '***' 0.001 '**' 0.01 '*' 0.05 '.' 0.1 ' ' 1
```

```
ado16S2 <- adonis(tcom16S ~ biol_16S$Flood_class * biol_16S$Sample, data = tcom16S, 
    type = "bray", permutations = 999, binary = FALSE)
ado16S2
```

```
## 
## Call:
## adonis(formula = tcom16S ~ biol_16S$Flood_class * biol_16S$Sample,      data = tcom16S, permutations = 999, type = "bray", binary = FALSE) 
## 
## Permutation: free
## Number of permutations: 999
## 
## Terms added sequentially (first to last)
## 
##                                      Df SumsOfSqs MeanSqs F.Model      R2
## biol_16S$Flood_class                  3    1.8697 0.62324  2.1230 0.18794
## biol_16S$Sample                       1    0.6232 0.62317  2.1228 0.06264
## biol_16S$Flood_class:biol_16S$Sample  3    0.9970 0.33234  1.1321 0.10022
## Residuals                            22    6.4584 0.29356         0.64920
## Total                                29    9.9483                 1.00000
##                                      Pr(>F)   
## biol_16S$Flood_class                  0.003 **
## biol_16S$Sample                       0.015 * 
## biol_16S$Flood_class:biol_16S$Sample  0.271   
## Residuals                                     
## Total                                         
## ---
## Signif. codes:  0 '***' 0.001 '**' 0.01 '*' 0.05 '.' 0.1 ' ' 1
```

```
# 18S

tcom18S <- t(Abu_VST_18S3)
tcom18S <- as.data.frame(tcom18S)


ado18S <- adonis2(tcom18S ~ biol_18S$Flood_class * biol_18S$Sample, data = tcom18S, 
    type = "bray", binary = FALSE, permutations = 999, by = NULL)
ado18S
```

```
## Permutation test for adonis under reduced model
## Permutation: free
## Number of permutations: 999
## 
## adonis2(formula = tcom18S ~ biol_18S$Flood_class * biol_18S$Sample, data = tcom18S, permutations = 999, by = NULL, type = "bray", binary = FALSE)
##          Df SumOfSqs      R2      F Pr(>F)  
## Model     7   2.5436 0.30978 1.3464  0.044 *
## Residual 21   5.6674 0.69022                
## Total    28   8.2109 1.00000                
## ---
## Signif. codes:  0 '***' 0.001 '**' 0.01 '*' 0.05 '.' 0.1 ' ' 1
```

```
ado18S2 <- adonis(tcom18S ~ biol_18S$Flood_class * biol_18S$Sample, data = tcom18S, 
    type = "bray", permutations = 999, binary = FALSE)
ado18S2
```

```
## 
## Call:
## adonis(formula = tcom18S ~ biol_18S$Flood_class * biol_18S$Sample,      data = tcom18S, permutations = 999, type = "bray", binary = FALSE) 
## 
## Permutation: free
## Number of permutations: 999
## 
## Terms added sequentially (first to last)
## 
##                                      Df SumsOfSqs MeanSqs F.Model      R2
## biol_18S$Flood_class                  3    1.2148 0.40493 1.50043 0.14795
## biol_18S$Sample                       1    0.6978 0.69785 2.58582 0.08499
## biol_18S$Flood_class:biol_18S$Sample  3    0.6309 0.21031 0.77927 0.07684
## Residuals                            21    5.6674 0.26988         0.69022
## Total                                28    8.2109                 1.00000
##                                      Pr(>F)   
## biol_18S$Flood_class                  0.049 * 
## biol_18S$Sample                       0.006 **
## biol_18S$Flood_class:biol_18S$Sample  0.846   
## Residuals                                     
## Total                                         
## ---
## Signif. codes:  0 '***' 0.001 '**' 0.01 '*' 0.05 '.' 0.1 ' ' 1
```

```
# 16S

sc16S2$Flood_class <- ordered(sc16S2$Flood_class, levels = c("H_Vz", "L_Vz", "M_Vz", 
    "TF"))

figNMDS_Shan_16S <- sc16S2 %>% ggplot(aes(x = NMDS1, y = NMDS2, colour = Flood_class, 
    shape = Sample)) + geom_point(position = position_jitter(0.1), size = 5) + # stat_ellipse(aes(fill=Flood_class), alpha=.05,type='t',size =1,
# geom='polygon')+
ggtitle("A) Below-ground prokaryotes (16S)") + xlab("") + ylab("") + scale_fill_manual(values = colsFlood) + 
    scale_color_manual(values = colsFlood) + theme(panel.grid.major = element_blank(), 
    panel.grid.minor = element_blank(), panel.background = element_blank(), axis.line = element_line(colour = "black"), 
    panel.border = element_rect(colour = "black", fill = NA, size = 1), legend.position = ("none"))

sc18S2$Flood_class <- ordered(sc18S2$Flood_class, levels = c("H_Vz", "L_Vz", "M_Vz", 
    "TF"))

figNMDS_Shan_18S <- sc18S2 %>% ggplot(aes(x = NMDS1, y = NMDS2, colour = Flood_class, 
    shape = Sample)) + geom_point(position = position_jitter(0.1), size = 5) + # stat_ellipse(aes(fill=Flood_class), alpha=.05,type='t',size =1,
# geom='polygon')+
ggtitle("B) Below-ground eukaryotes (18S)") + xlab("") + ylab("") + scale_fill_manual(values = colsFlood) + 
    scale_color_manual(values = colsFlood) + theme(panel.grid.major = element_blank(), 
    panel.grid.minor = element_blank(), panel.background = element_blank(), axis.line = element_line(colour = "black"), 
    panel.border = element_rect(colour = "black", fill = NA, size = 1), legend.position = ("none"))

figNMDSPlants <- plants2 %>% ggplot(aes(x = NMDS1, y = NMDS2, colour = Flood_class)) + 
    geom_point(position = position_jitter(0.1), shape = 15, size = 5) + # stat_ellipse(aes(fill=Flood_class), alpha=.05,type='t',size =1,
# geom='polygon')+
ggtitle("C) Above-ground woody plant") + xlab("") + ylab("") + scale_fill_manual(values = colsFlood) + 
    scale_color_manual(values = colsFlood) + theme(panel.grid.major = element_blank(), 
    panel.grid.minor = element_blank(), panel.background = element_blank(), axis.line = element_line(colour = "black"), 
    panel.border = element_rect(colour = "black", fill = NA, size = 1), legend.position = ("none"))

figNMDSShan <- ggarrange(figNMDS_Shan_16S, figNMDS_Shan_18S, figNMDSPlants, common.legend = TRUE, 
    legend = "bottom", ncol = 3)
ggsave("figNMDSShan.pdf", plot = figNMDSShan, width = 12, height = 8)
figNMDSShan
```

```
############################################################ 16S load and prepare data taxonomy
tax_16S <- read.csv("input/16S_taxonomy_clean.csv")
tax_16S <- tax_16S %>% dplyr::select(ASV_ID = ASV_ID, taxonomy = classifications)

# View(tax_16S)
head(tax_16S)
```

```
##     ASV_ID
## 1 ASV10000
## 2 ASV10007
## 3  ASV1000
## 4 ASV10012
## 5 ASV10017
## 6 ASV10023
##                                                                                                     taxonomy
## 1      silva|138|43606|Bacteria;Actinobacteriota;Actinobacteria;Micromonosporales;Micromonosporaceae;Asanoa;
## 2 silva|138|43354|Bacteria;Actinobacteriota;Actinobacteria;Corynebacteriales;Mycobacteriaceae;Mycobacterium;
## 3                                                                silva|119|1448|Bacteria;Chloroflexi;KD4-96;
## 4  silva|138|43300|Bacteria;Actinobacteriota;Acidimicrobiia;Microtrichales;Ilumatobacteraceae;Ilumatobacter;
## 5  silva|132|26244|Bacteria;Proteobacteria;Alphaproteobacteria;Sphingomonadales;Sphingomonadaceae;Ellin6055;
## 6          silva|138|43378|Bacteria;Actinobacteriota;Actinobacteria;Frankiales;Acidothermaceae;Acidothermus;
```

```
tail(tax_16S)
```

```
##        ASV_ID
## 10205 ASV9967
## 10206 ASV9973
## 10207  ASV998
## 10208 ASV9981
## 10209 ASV9984
## 10210 ASV9997
##                                                                                                            taxonomy
## 10205    silva|138|43648|Bacteria;Actinobacteriota;Actinobacteria;Propionibacteriales;Nocardioidaceae;Nocardioides;
## 10206          silva|138|43412|Bacteria;Actinobacteriota;Actinobacteria;Kineosporiales;Kineosporiaceae;Kineosporia;
## 10207                                           silva|138|43211|Bacteria;Acidobacteriota;Acidobacteriae;Subgroup 2;
## 10208 silva|119|2392|Bacteria;Proteobacteria;Alphaproteobacteria;Caulobacterales;Caulobacteraceae;Phenylobacterium;
## 10209  silva|138|43688|Bacteria;Actinobacteriota;Actinobacteria;Pseudonocardiales;Pseudonocardiaceae;Amycolatopsis;
## 10210 silva|138|43616|Bacteria;Actinobacteriota;Actinobacteria;Micromonosporales;Micromonosporaceae;Micromonospora;
```

```
### abundance per site

abus_16S <- read_csv("input/ASV_table_16S.csv")
```

```
## Parsed with column specification:
## cols(
##   .default = col_double(),
##   ASV_ID = col_character()
## )
```

```
## See spec(...) for full column specifications.
```

```
head(abus_16S)
```

```
## Warning: `...` is not empty.
## 
## We detected these problematic arguments:
## * `needs_dots`
## 
## These dots only exist to allow future extensions and should be empty.
## Did you misspecify an argument?
```

```
## # A tibble: 6 x 31
##   ASV_ID TF12S TF17S LV15S LV14S LV16S  MV2S  HV5S  HV8S  HV7S TF19S  TF9S LV10S
##   <chr>  <dbl> <dbl> <dbl> <dbl> <dbl> <dbl> <dbl> <dbl> <dbl> <dbl> <dbl> <dbl>
## 1 ASV91~     2     0     0     0     0     0     0     0     0     0     0     0
## 2 ASV91~     2     0     0     0     0     0     0     0     0     0     0     0
## 3 ASV91~     2     0     0     0     0     0     0     0     0     0     0     0
## 4 ASV91~     2     0     0     0     0     0     0     0     0     0     0     0
## 5 ASV91~     1     0     0     0     1     0     0     0     0     0     0     0
## 6 ASV91~     1     0     0     0     0     0     0     0     0     0     0     0
## # ... with 18 more variables: LV13S <dbl>, TF12L <dbl>, TF17L <dbl>,
## #   LV15L <dbl>, LV14L <dbl>, LV16L <dbl>, MV6L <dbl>, MV3L <dbl>, HV5L <dbl>,
## #   LV4L <dbl>, HV8L <dbl>, MV18L <dbl>, TF19L <dbl>, TF9L <dbl>, MV11L <dbl>,
## #   LV10L <dbl>, LV13L <dbl>, HV1L <dbl>
```

```
nams_16S <- abus_16S$ASV_ID

abus_16S$ASV_ID <- nams_16S


## merge taxonomy and abundance
abutaxs_16S <- inner_join(abus_16S, tax_16S, by = "ASV_ID")
head(abutaxs_16S)
```

```
## Warning: `...` is not empty.
## 
## We detected these problematic arguments:
## * `needs_dots`
## 
## These dots only exist to allow future extensions and should be empty.
## Did you misspecify an argument?
```

```
## # A tibble: 6 x 32
##   ASV_ID TF12S TF17S LV15S LV14S LV16S  MV2S  HV5S  HV8S  HV7S TF19S  TF9S LV10S
##   <chr>  <dbl> <dbl> <dbl> <dbl> <dbl> <dbl> <dbl> <dbl> <dbl> <dbl> <dbl> <dbl>
## 1 ASV91~     2     0     0     0     0     0     0     0     0     0     0     0
## 2 ASV91~     2     0     0     0     0     0     0     0     0     0     0     0
## 3 ASV91~     2     0     0     0     0     0     0     0     0     0     0     0
## 4 ASV91~     2     0     0     0     0     0     0     0     0     0     0     0
## 5 ASV91~     1     0     0     0     1     0     0     0     0     0     0     0
## 6 ASV91~     1     0     0     0     0     0     0     0     0     0     0     0
## # ... with 19 more variables: LV13S <dbl>, TF12L <dbl>, TF17L <dbl>,
## #   LV15L <dbl>, LV14L <dbl>, LV16L <dbl>, MV6L <dbl>, MV3L <dbl>, HV5L <dbl>,
## #   LV4L <dbl>, HV8L <dbl>, MV18L <dbl>, TF19L <dbl>, TF9L <dbl>, MV11L <dbl>,
## #   LV10L <dbl>, LV13L <dbl>, HV1L <dbl>, taxonomy <chr>
```

```
write.csv(abutaxs_16S, "input/16S_taxonomy_metadata_16S2.csv")

### Identity and number of ASVs not in the taxonomy file
sum(is.na(abutaxs_16S$taxonomy))
```

```
## [1] 0
```

```
sum(is.na(abutaxs_16S$taxonomy))/nrow(abus_16S)
```

```
## [1] 0
```

```
### Identity and number of ASVs not in the taxonomy file
sum(is.na(abutaxs_16S$STF2))
```

```
## Warning: Unknown or uninitialised column: `STF2`.
```

```
## [1] 0
```

```
sum(is.na(abutaxs_16S$STF2))/nrow(tax_16S)
```

```
## Warning: Unknown or uninitialised column: `STF2`.
```

```
## [1] 0
```

```
## Create taxonomy table not rarefied put table in right format for summary
snon_rar_16S <- gather(abutaxs_16S, key = site, value = presence, -ASV_ID, -taxonomy) %>% 
    filter(presence > 0)


#### define the taxonomic level to use for all groups,and add to table
tax.list <- c("Actinobacteria", "Acidimicrobiia", "Alphaproteobacteria", "Acidobacteriota", 
    "Chloroflexi", "Planctomycetota", "Patescibacteria")


snon_rar_16S <- snon_rar_16S %>% mutate(tax.class = "others") %>% mutate(tax.class = ifelse(grepl(tax.list[1], 
    taxonomy), tax.list[1], tax.class)) %>% mutate(tax.class = ifelse(grepl(tax.list[2], 
    taxonomy), tax.list[2], tax.class)) %>% mutate(tax.class = ifelse(grepl(tax.list[3], 
    taxonomy), tax.list[3], tax.class)) %>% mutate(tax.class = ifelse(grepl(tax.list[4], 
    taxonomy), tax.list[4], tax.class)) %>% mutate(tax.class = ifelse(grepl(tax.list[5], 
    taxonomy), tax.list[5], tax.class)) %>% mutate(tax.class = ifelse(grepl(tax.list[6], 
    taxonomy), tax.list[6], tax.class)) %>% mutate(tax.class = ifelse(grepl(tax.list[7], 
    taxonomy), tax.list[7], tax.class)) %>% mutate(tax.class = ifelse(is.na(taxonomy), 
    "unknown", tax.class))


#### create output tables
head(abus_16S)
```

```
## Warning: `...` is not empty.
## 
## We detected these problematic arguments:
## * `needs_dots`
## 
## These dots only exist to allow future extensions and should be empty.
## Did you misspecify an argument?
```

```
## # A tibble: 6 x 31
##   ASV_ID TF12S TF17S LV15S LV14S LV16S  MV2S  HV5S  HV8S  HV7S TF19S  TF9S LV10S
##   <chr>  <dbl> <dbl> <dbl> <dbl> <dbl> <dbl> <dbl> <dbl> <dbl> <dbl> <dbl> <dbl>
## 1 ASV91~     2     0     0     0     0     0     0     0     0     0     0     0
## 2 ASV91~     2     0     0     0     0     0     0     0     0     0     0     0
## 3 ASV91~     2     0     0     0     0     0     0     0     0     0     0     0
## 4 ASV91~     2     0     0     0     0     0     0     0     0     0     0     0
## 5 ASV91~     1     0     0     0     1     0     0     0     0     0     0     0
## 6 ASV91~     1     0     0     0     0     0     0     0     0     0     0     0
## # ... with 18 more variables: LV13S <dbl>, TF12L <dbl>, TF17L <dbl>,
## #   LV15L <dbl>, LV14L <dbl>, LV16L <dbl>, MV6L <dbl>, MV3L <dbl>, HV5L <dbl>,
## #   LV4L <dbl>, HV8L <dbl>, MV18L <dbl>, TF19L <dbl>, TF9L <dbl>, MV11L <dbl>,
## #   LV10L <dbl>, LV13L <dbl>, HV1L <dbl>
```

```
# create site to habitat and sample key
skeys_16S <- names(abus_16S) %>% as_tibble() %>% dplyr::rename(site = "value") %>% 
    mutate(habitat = "TF") %>% mutate(habitat = ifelse(grepl("LV", site), "LV", habitat)) %>% 
    mutate(habitat = ifelse(grepl("MV", site), "MV", habitat)) %>% mutate(habitat = ifelse(grepl("HV", 
    site), "HV", habitat)) %>% mutate(sample = "In") %>% mutate(sample = ifelse(grepl("L", 
    site), "L", sample)) %>% mutate(sample = ifelse(grepl("S", site), "S", sample))


out_abus_16S <- snon_rar_16S %>% left_join(skeys_16S, by = "site")

tail(out_abus_16S)
```

```
## Warning: `...` is not empty.
## 
## We detected these problematic arguments:
## * `needs_dots`
## 
## These dots only exist to allow future extensions and should be empty.
## Did you misspecify an argument?
```

```
## # A tibble: 6 x 7
##   ASV_ID taxonomy                      site  presence tax.class   habitat sample
##   <chr>  <chr>                         <chr>    <dbl> <chr>       <chr>   <chr> 
## 1 ASV6   silva|132|26151|Bacteria;Pro~ HV1L        29 Alphaprote~ HV      L     
## 2 ASV5   silva|132|26151|Bacteria;Pro~ HV1L         6 Alphaprote~ HV      L     
## 3 ASV4   silva|119|2554|Bacteria;Prot~ HV1L         4 Alphaprote~ HV      L     
## 4 ASV3   silva|132|26151|Bacteria;Pro~ HV1L        18 Alphaprote~ HV      L     
## 5 ASV2   silva|138|43378|Bacteria;Act~ HV1L         2 Actinobact~ HV      L     
## 6 ASV1   silva|132|26151|Bacteria;Pro~ HV1L       188 Alphaprote~ HV      L
```

```
write_csv(out_abus_16S, "16S_taxonomy_per_site.csv")

############################################################ 18S load and prepare data taxonomy
tax_18S <- read.csv("input/18S_taxonomy_clean.csv")
tax_18S <- tax_18S %>% dplyr::select(ASV_ID = ASV_ID, taxonomy = classifications)

# View(tax_18S)
head(tax_18S)
```

```
##    ASV_ID
## 1    ASV1
## 2   ASV10
## 3 ASV1005
## 4 ASV1012
## 5 ASV1019
## 6 ASV1026
##                                                                                                                                                taxonomy
## 1      Eukaryota;Amorphea;Obazoa;Opisthokonta;Nucletmycea;Fungi;Dikarya;Ascomycota;Pezizomycotina;Eurotiomycetes;Eurotiales;Aspergillaceae;Aspergillus;
## 2 Eukaryota;Amorphea;Obazoa;Opisthokonta;Nucletmycea;Fungi;Dikarya;Ascomycota;Pezizomycotina;Dothideomycetes;Capnodiales;Mycosphaerellaceae;Cercospora;
## 3      Eukaryota;Amorphea;Obazoa;Opisthokonta;Nucletmycea;Fungi;Dikarya;Ascomycota;Pezizomycotina;Eurotiomycetes;Eurotiales;Trichocomaceae;Thermomyces;
## 4                                                                                Eukaryota;Amorphea;Amoebozoa;Discosea;Flabellinia;Vannellida;Vannella;
## 5                 Eukaryota;Amorphea;Obazoa;Opisthokonta;Nucletmycea;Fungi;Dikarya;Ascomycota;Pezizomycotina;Eurotiomycetes;Chaetothyriales;uncultured;
## 6                                                                                                                         Eukaryota;Amorphea;Amoebozoa;
```

```
tail(tax_18S)
```

```
##      ASV_ID
## 2047 ASV963
## 2048 ASV970
## 2049 ASV978
## 2050 ASV984
## 2051 ASV990
## 2052 ASV997
##                                                                                                                                                   taxonomy
## 2047                                                                                                               Eukaryota;Amorphea;Amoebozoa;Mycamoeba;
## 2048 Eukaryota;Amorphea;Obazoa;Opisthokonta;Nucletmycea;Fungi;Dikarya;Basidiomycota;Agaricomycotina;Agaricomycetes;Corticiales;Corticiaceae;Peniophorella;
## 2049                                             Eukaryota;SAR;Alveolata;Ciliophora;Intramacronucleata;Conthreep;Oligohymenophorea;Peritrichia;Vorticella;
## 2050                                                                                            Eukaryota;Amorphea;Amoebozoa;Tubulinea;Euamoebida;BOLA868;
## 2051   Eukaryota;Amorphea;Obazoa;Opisthokonta;Nucletmycea;Fungi;Dikarya;Basidiomycota;Agaricomycotina;Agaricomycetes;Agaricales;Agaricaceae;Leucoagaricus;
## 2052                                                             Eukaryota;SAR;Rhizaria;Cercozoa;Imbricatea;Silicofilosea;Euglyphida;Trinematidae;Trinema;
```

```
### abundance per site

abus_18S <- read_csv("input/ASV_table_18S.csv")
```

```
## Parsed with column specification:
## cols(
##   .default = col_double(),
##   ASV_ID = col_character()
## )
## See spec(...) for full column specifications.
```

```
head(abus_18S)
```

```
## Warning: `...` is not empty.
## 
## We detected these problematic arguments:
## * `needs_dots`
## 
## These dots only exist to allow future extensions and should be empty.
## Did you misspecify an argument?
```

```
## # A tibble: 6 x 30
##   ASV_ID TF12S TF17S LV15S LV14S LV16S  MV2S  HV5S  HV8S  HV7S TF19S  TF9S LV10S
##   <chr>  <dbl> <dbl> <dbl> <dbl> <dbl> <dbl> <dbl> <dbl> <dbl> <dbl> <dbl> <dbl>
## 1 ASV10~     5     0     0     0     0     1     0     0     0     0     0     1
## 2 ASV12~     1     0     0     0     0     0     0     0     0     0     0     0
## 3 ASV15~     1     0     0     0     0     0     0     0     0     0     0     0
## 4 ASV19~     0     0     2     0     0     2     0     0     0     0     0     0
## 5 ASV19~     0     0     0     0     0     0     0     0     0     0     0     0
## 6 ASV21~     0     0     0     0     0     0     0     0     0     0     0     0
## # ... with 17 more variables: LV13S <dbl>, TF12L <dbl>, TF17L <dbl>,
## #   LV15L <dbl>, LV14L <dbl>, LV16L <dbl>, MV6L <dbl>, MV3L <dbl>, HV5L <dbl>,
## #   LV4L <dbl>, HV8L <dbl>, MV18L <dbl>, TF19L <dbl>, TF9L <dbl>, LV10L <dbl>,
## #   LV13L <dbl>, HV1L <dbl>
```

```
nams_18S <- abus_18S$ASV_ID

abus_18S$ASV_ID <- nams_18S


## merge taxonomy and abundance
abutaxs_18S <- inner_join(abus_18S, tax_18S, by = "ASV_ID")
head(abutaxs_18S)
```

```
## Warning: `...` is not empty.
## 
## We detected these problematic arguments:
## * `needs_dots`
## 
## These dots only exist to allow future extensions and should be empty.
## Did you misspecify an argument?
```

```
## # A tibble: 6 x 31
##   ASV_ID TF12S TF17S LV15S LV14S LV16S  MV2S  HV5S  HV8S  HV7S TF19S  TF9S LV10S
##   <chr>  <dbl> <dbl> <dbl> <dbl> <dbl> <dbl> <dbl> <dbl> <dbl> <dbl> <dbl> <dbl>
## 1 ASV10~     5     0     0     0     0     1     0     0     0     0     0     1
## 2 ASV12~     1     0     0     0     0     0     0     0     0     0     0     0
## 3 ASV15~     1     0     0     0     0     0     0     0     0     0     0     0
## 4 ASV19~     0     0     0     0     0     0     0     0     0     0     0     0
## 5 ASV765     0     0     0     0     0     0     0     0     0     0     0     0
## 6 ASV580     1     0     0     0     0     0     0     0     0     0     0     0
## # ... with 18 more variables: LV13S <dbl>, TF12L <dbl>, TF17L <dbl>,
## #   LV15L <dbl>, LV14L <dbl>, LV16L <dbl>, MV6L <dbl>, MV3L <dbl>, HV5L <dbl>,
## #   LV4L <dbl>, HV8L <dbl>, MV18L <dbl>, TF19L <dbl>, TF9L <dbl>, LV10L <dbl>,
## #   LV13L <dbl>, HV1L <dbl>, taxonomy <chr>
```

```
write.csv(abutaxs_18S, "input/18S_taxonomy_metadata_18S2.csv")

### Identity and number of ASVs not in the taxonomy file
sum(is.na(abutaxs_18S$taxonomy))
```

```
## [1] 0
```

```
sum(is.na(abutaxs_18S$taxonomy))/nrow(abus_18S)
```

```
## [1] 0
```

```
### Identity and number of ASVs not in the taxonomy file
sum(is.na(abutaxs_18S$STF2))
```

```
## Warning: Unknown or uninitialised column: `STF2`.
```

```
## [1] 0
```

```
sum(is.na(abutaxs_18S$STF2))/nrow(tax_18S)
```

```
## Warning: Unknown or uninitialised column: `STF2`.
```

```
## [1] 0
```

```
## Create taxonomy table not rarefied put table in right format for summary
snon_rar_18S <- gather(abutaxs_18S, key = site, value = presence, -ASV_ID, -taxonomy) %>% 
    filter(presence > 0)


#### define the taxonomic level to use for all groups,and add to table
tax.list <- c("Fungi", "Amoebozoa", "Ciliophora", "Conoidasida", "Dinoflagellata", 
    "Protalveolata", "Cercozoa", "Stramenopiles", "Chloroplastida", "Choanozoa")


snon_rar_18S <- snon_rar_18S %>% mutate(tax.class = "others") %>% mutate(tax.class = ifelse(grepl(tax.list[1], 
    taxonomy), tax.list[1], tax.class)) %>% mutate(tax.class = ifelse(grepl(tax.list[2], 
    taxonomy), tax.list[2], tax.class)) %>% mutate(tax.class = ifelse(grepl(tax.list[3], 
    taxonomy), tax.list[3], tax.class)) %>% mutate(tax.class = ifelse(grepl(tax.list[4], 
    taxonomy), tax.list[4], tax.class)) %>% mutate(tax.class = ifelse(grepl(tax.list[5], 
    taxonomy), tax.list[5], tax.class)) %>% mutate(tax.class = ifelse(grepl(tax.list[6], 
    taxonomy), tax.list[6], tax.class)) %>% mutate(tax.class = ifelse(grepl(tax.list[7], 
    taxonomy), tax.list[7], tax.class)) %>% mutate(tax.class = ifelse(grepl(tax.list[8], 
    taxonomy), tax.list[8], tax.class)) %>% mutate(tax.class = ifelse(grepl(tax.list[9], 
    taxonomy), tax.list[9], tax.class)) %>% mutate(tax.class = ifelse(grepl(tax.list[10], 
    taxonomy), tax.list[10], tax.class)) %>% mutate(tax.class = ifelse(is.na(taxonomy), 
    "unknown", tax.class))


#### create output tables
head(abus_18S)
```

```
## Warning: `...` is not empty.
## 
## We detected these problematic arguments:
## * `needs_dots`
## 
## These dots only exist to allow future extensions and should be empty.
## Did you misspecify an argument?
```

```
## # A tibble: 6 x 30
##   ASV_ID TF12S TF17S LV15S LV14S LV16S  MV2S  HV5S  HV8S  HV7S TF19S  TF9S LV10S
##   <chr>  <dbl> <dbl> <dbl> <dbl> <dbl> <dbl> <dbl> <dbl> <dbl> <dbl> <dbl> <dbl>
## 1 ASV10~     5     0     0     0     0     1     0     0     0     0     0     1
## 2 ASV12~     1     0     0     0     0     0     0     0     0     0     0     0
## 3 ASV15~     1     0     0     0     0     0     0     0     0     0     0     0
## 4 ASV19~     0     0     2     0     0     2     0     0     0     0     0     0
## 5 ASV19~     0     0     0     0     0     0     0     0     0     0     0     0
## 6 ASV21~     0     0     0     0     0     0     0     0     0     0     0     0
## # ... with 17 more variables: LV13S <dbl>, TF12L <dbl>, TF17L <dbl>,
## #   LV15L <dbl>, LV14L <dbl>, LV16L <dbl>, MV6L <dbl>, MV3L <dbl>, HV5L <dbl>,
## #   LV4L <dbl>, HV8L <dbl>, MV18L <dbl>, TF19L <dbl>, TF9L <dbl>, LV10L <dbl>,
## #   LV13L <dbl>, HV1L <dbl>
```

```
# create site to habitat and sample key
skeys_18S <- names(abus_18S) %>% as_tibble() %>% dplyr::rename(site = "value") %>% 
    mutate(habitat = "TF") %>% mutate(habitat = ifelse(grepl("LV", site), "LV", habitat)) %>% 
    mutate(habitat = ifelse(grepl("MV", site), "MV", habitat)) %>% mutate(habitat = ifelse(grepl("HV", 
    site), "HV", habitat)) %>% mutate(sample = "In") %>% mutate(sample = ifelse(grepl("L", 
    site), "L", sample)) %>% mutate(sample = ifelse(grepl("S", site), "S", sample))


out_abus_18S <- snon_rar_18S %>% left_join(skeys_18S, by = "site")

tail(out_abus_18S)
```

```
## Warning: `...` is not empty.
## 
## We detected these problematic arguments:
## * `needs_dots`
## 
## These dots only exist to allow future extensions and should be empty.
## Did you misspecify an argument?
```

```
## # A tibble: 6 x 7
##   ASV_ID taxonomy                        site  presence tax.class habitat sample
##   <chr>  <chr>                           <chr>    <dbl> <chr>     <chr>   <chr> 
## 1 ASV949 Eukaryota;SAR;Alveolata;Ciliop~ HV1L         4 Ciliopho~ HV      L     
## 2 ASV967 Eukaryota;SAR;Alveolata;Apicom~ HV1L         2 Conoidas~ HV      L     
## 3 ASV976 Eukaryota;SAR;Rhizaria;Cercozo~ HV1L         6 Cercozoa  HV      L     
## 4 ASV98  Eukaryota;Amorphea;Obazoa;Opis~ HV1L         1 Fungi     HV      L     
## 5 ASV998 Eukaryota;Amorphea;Obazoa;Opis~ HV1L        10 Choanozoa HV      L     
## 6 ASV999 Eukaryota;SAR;Rhizaria;Cercozo~ HV1L         2 Cercozoa  HV      L
```

```
write_csv(out_abus_18S, "18S_taxonomy_per_site.csv")
```

```
###################################################################### Plots taxonomic groups############################

# 16S

head(out_abus_16S)
```

```
## Warning: `...` is not empty.
## 
## We detected these problematic arguments:
## * `needs_dots`
## 
## These dots only exist to allow future extensions and should be empty.
## Did you misspecify an argument?
```

```
## # A tibble: 6 x 7
##   ASV_ID  taxonomy                     site  presence tax.class   habitat sample
##   <chr>   <chr>                        <chr>    <dbl> <chr>       <chr>   <chr> 
## 1 ASV9119 silva|138|43211|Bacteria;Ac~ TF12S        2 Acidobacte~ TF      S     
## 2 ASV9120 silva|138|43695|Bacteria;Ac~ TF12S        2 Actinobact~ TF      S     
## 3 ASV9121 silva|132|25197|Bacteria;Ch~ TF12S        2 Chloroflexi TF      S     
## 4 ASV9122 silva|119|1389|Bacteria;Chl~ TF12S        2 Chloroflexi TF      S     
## 5 ASV9123 silva|132|26132|Bacteria;Pr~ TF12S        1 Alphaprote~ TF      S     
## 6 ASV9124 silva|138|43378|Bacteria;Ac~ TF12S        1 Actinobact~ TF      S
```

```
# load 16S
hab_16S <- out_abus_16S %>% distinct(ASV_ID, habitat, .keep_all = T) %>% group_by(habitat, 
    tax.class) %>% summarise(ASVs.number = n()) %>% mutate(fraction.ASVs = round(ASVs.number/sum(ASVs.number), 
    3)) %>% filter(!is.na(habitat)) %>% dplyr::rename(value = habitat) %>% mutate(type = "habitat")
```

```
## `summarise()` regrouping output by 'habitat' (override with `.groups` argument)
```

```
ord <- c("TF", "HV", "MV", "LV")
hab_16S$value <- factor(hab_16S$value, levels = ord)

ord2 <- c("Alphaproteobacteria", "Actinobacteria", "Acidimicrobiia", "Acidobacteriota", 
    "Chloroflexi", "Planctomycetota", "Patescibacteria", "others", "unknown")
hab_16S$tax.class <- factor(hab_16S$tax.class, levels = ord2)


# 18S

head(out_abus_18S)
```

```
## Warning: `...` is not empty.
## 
## We detected these problematic arguments:
## * `needs_dots`
## 
## These dots only exist to allow future extensions and should be empty.
## Did you misspecify an argument?
```

```
## # A tibble: 6 x 7
##   ASV_ID  taxonomy                      site  presence tax.class  habitat sample
##   <chr>   <chr>                         <chr>    <dbl> <chr>      <chr>   <chr> 
## 1 ASV1094 Eukaryota;Amorphea;Obazoa;Op~ TF12S        5 Fungi      TF      S     
## 2 ASV1250 Eukaryota;SAR;Alveolata;Cili~ TF12S        1 Ciliophora TF      S     
## 3 ASV1599 Eukaryota;SAR;Alveolata;Cili~ TF12S        1 Ciliophora TF      S     
## 4 ASV580  Eukaryota;Archaeplastida;Chl~ TF12S        1 Chloropla~ TF      S     
## 5 ASV50   Eukaryota;Amorphea;Obazoa;Op~ TF12S       16 Fungi      TF      S     
## 6 ASV850  Eukaryota;Amorphea;Obazoa;Op~ TF12S        1 Fungi      TF      S
```

```
# load 18S
hab_18S <- out_abus_18S %>% distinct(ASV_ID, habitat, .keep_all = T) %>% group_by(habitat, 
    tax.class) %>% summarise(ASVs.number = n()) %>% mutate(fraction.ASVs = round(ASVs.number/sum(ASVs.number), 
    3)) %>% filter(!is.na(habitat)) %>% dplyr::rename(value = habitat) %>% mutate(type = "habitat")
```

```
## `summarise()` regrouping output by 'habitat' (override with `.groups` argument)
```

```
ord <- c("TF", "HV", "MV", "LV")
hab_18S$value <- factor(hab_18S$value, levels = ord)

ord2 <- c("Fungi", "Chloroplastida", "Choanozoa", "Amoebozoa", "Cercozoa", "Ciliophora", 
    "Conoidasida", "Dinoflagellata", "Protalveolata", "Stramenopiles", "others", 
    "unknown")
hab_18S$tax.class <- factor(hab_18S$tax.class, levels = ord2)


# number

cols <- c("#00441b", "#238b45", "#74c476", "#c7e9c0")

Hab_N_16S_plot <- ggplot(data = hab_16S, aes(x = tax.class, y = ASVs.number, fill = value)) + 
    geom_bar(stat = "identity", position = position_dodge2()) + scale_fill_manual(values = cols, 
    name = "") + ggtitle("A) Prokaryotic taxonomic profile (16S)") + ylab("Number of ASVs") + 
    xlab("") + coord_flip() + theme(panel.grid.major = element_blank(), panel.grid.minor = element_blank(), 
    panel.background = element_blank(), axis.line = element_line(colour = "black"), 
    panel.border = element_rect(colour = "black", fill = NA, size = 1), legend.position = "none")


Hab_N_18S_plot <- ggplot(data = hab_18S, aes(x = tax.class, y = ASVs.number, fill = value)) + 
    geom_bar(stat = "identity", position = position_dodge2()) + scale_fill_manual(values = cols, 
    name = "") + ggtitle("B) Eukaryotic taxonomic profile (18S)") + ylab("B) Number of ASVs") + 
    xlab("") + coord_flip() + theme(panel.grid.major = element_blank(), panel.grid.minor = element_blank(), 
    panel.background = element_blank(), axis.line = element_line(colour = "black"), 
    panel.border = element_rect(colour = "black", fill = NA, size = 1), legend.position = "none")

figTaxNumber <- ggarrange(Hab_N_16S_plot, Hab_N_18S_plot, common.legend = TRUE, legend = "bottom", 
    ncol = 2)
ggsave("figTaxNumber.pdf", plot = figTaxNumber, width = 12, height = 8)
figTaxNumber
```

```
# fraction

Hab_F_16S_plot <- ggplot(data = hab_16S, aes(x = tax.class, y = fraction.ASVs, fill = value)) + 
    geom_bar(stat = "identity", position = position_dodge()) + scale_fill_manual(values = cols, 
    name = "") + ggtitle("A) Prokaryotic taxonomic profile (16S)") + ylab("Fraction of ASVs") + 
    xlab("") + coord_flip() + theme_bw() + theme(panel.grid.major.x = element_line(size = 0.5, 
    colour = "#d9d9d9"), panel.grid.major.y = element_blank(), panel.grid.minor.x = element_line(size = 0.5, 
    colour = "#f0f0f0"), panel.grid.minor.y = element_blank(), panel.background = element_blank(), 
    axis.line = element_line(colour = "black"), panel.border = element_rect(colour = "black", 
        fill = NA, size = 1), legend.position = "right")


Hab_F_18S_plot <- ggplot(data = hab_18S, aes(x = tax.class, y = fraction.ASVs, fill = value)) + 
    geom_bar(stat = "identity", position = position_dodge()) + scale_fill_manual(values = cols, 
    name = "") + ggtitle("B) Eukaryotic taxonomic profile (18S)") + ylab("Fraction of ASVs") + 
    xlab("") + coord_flip() + theme_bw() + theme(panel.grid.major.x = element_line(size = 0.5, 
    colour = "#d9d9d9"), panel.grid.major.y = element_blank(), panel.grid.minor.x = element_line(size = 0.5, 
    colour = "#f0f0f0"), panel.grid.minor.y = element_blank(), panel.background = element_blank(), 
    axis.line = element_line(colour = "black"), panel.border = element_rect(colour = "black", 
        fill = NA, size = 1), legend.position = "right")

figTaxFraction <- ggarrange(Hab_F_16S_plot, Hab_F_18S_plot, common.legend = TRUE, 
    legend = "bottom", ncol = 2)
ggsave("figTaxFraction.pdf", plot = figTaxFraction, width = 12, height = 8)
figTaxFraction
```

```
######################################### ASV 'ranges'

head(out_abus_16S)
```

```
## Warning: `...` is not empty.
## 
## We detected these problematic arguments:
## * `needs_dots`
## 
## These dots only exist to allow future extensions and should be empty.
## Did you misspecify an argument?
```

```
## # A tibble: 6 x 7
##   ASV_ID  taxonomy                     site  presence tax.class   habitat sample
##   <chr>   <chr>                        <chr>    <dbl> <chr>       <chr>   <chr> 
## 1 ASV9119 silva|138|43211|Bacteria;Ac~ TF12S        2 Acidobacte~ TF      S     
## 2 ASV9120 silva|138|43695|Bacteria;Ac~ TF12S        2 Actinobact~ TF      S     
## 3 ASV9121 silva|132|25197|Bacteria;Ch~ TF12S        2 Chloroflexi TF      S     
## 4 ASV9122 silva|119|1389|Bacteria;Chl~ TF12S        2 Chloroflexi TF      S     
## 5 ASV9123 silva|132|26132|Bacteria;Pr~ TF12S        1 Alphaprote~ TF      S     
## 6 ASV9124 silva|138|43378|Bacteria;Ac~ TF12S        1 Actinobact~ TF      S
```

```
pn_16S <- out_abus_16S %>% select(-taxonomy, -habitat) %>% group_by(tax.class, ASV_ID) %>% 
    summarise(plot_number = n()) %>% ungroup()
```

```
## `summarise()` regrouping output by 'tax.class' (override with `.groups` argument)
```

```
# number of habitats
hab_n_16S <- out_abus_16S %>% filter(!is.na(habitat)) %>% select(-taxonomy) %>% group_by(tax.class, 
    ASV_ID, habitat) %>% summarise(ASVs.number = n()) %>% ungroup() %>% select(-ASVs.number) %>% 
    group_by(tax.class, ASV_ID) %>% summarise(habitat_number = n()) %>% group_by(tax.class, 
    habitat_number) %>% summarise(num_hab_num = n()) %>% mutate(fraction_hab_num = round(num_hab_num/sum(num_hab_num), 
    3)) %>% ungroup()
```

```
## `summarise()` regrouping output by 'tax.class', 'ASV_ID' (override with `.groups` argument)
## `summarise()` regrouping output by 'tax.class' (override with `.groups` argument)
## `summarise()` regrouping output by 'tax.class' (override with `.groups` argument)
```

```
cols_tax <- c("#8dd3c7", "#ffffb3", "#bebada", "#fb8072", "#80b1d3", "#fdb462", "#b3de69", 
    "#fccde5", "#d9d9d9", "#bc80bd", "#ccebc5")

plo_pn_16S <- ggplot(data = pn_16S, aes(x = tax.class, y = plot_number, fill = tax.class)) + 
    geom_bar(stat = "identity", position = "dodge") + scale_fill_manual(values = cols_tax, 
    name = "") + ylab("Number of plots") + ggtitle("A) Number of plots") + theme_bw() + 
    theme(legend.position = "none") + theme(panel.grid.major = element_blank(), panel.grid.minor = element_blank(), 
    panel.background = element_blank(), axis.line = element_line(colour = "black"), 
    panel.border = element_rect(colour = "black", fill = NA, size = 1))

plo_pn_den_16S <- ggplot() + geom_density(data = pn_16S, aes(plot_number, group = tax.class, 
    col = tax.class), size = 1) + scale_colour_manual(values = cols_tax, name = "") + 
    ggtitle("B) Number of plots - density") + xlab("Number of plots") + ylab("Density") + 
    xlim(c(1, 39)) + guides(color = guide_legend(ncol = 2)) + theme_bw() + theme(panel.grid.major = element_blank(), 
    panel.grid.minor = element_blank(), panel.background = element_blank(), axis.line = element_line(colour = "black"), 
    panel.border = element_rect(colour = "black", fill = NA, size = 1), legend.position = c(0.65, 
        0.78), legend.title = element_blank())


plo_habn_16S <- ggplot(data = hab_n_16S, aes(x = tax.class, y = fraction_hab_num, 
    fill = as.factor(habitat_number))) + geom_bar(stat = "identity", position = position_dodge()) + 
    scale_fill_brewer(palette = "Blues", name = "") + ylab("Fraction of ASVs") + 
    ggtitle("C) Number of Flood Class") + guides(fill = guide_legend(ncol = 4)) + 
    theme_bw() + theme(panel.grid.major = element_blank(), panel.grid.minor = element_blank(), 
    panel.background = element_blank(), axis.line = element_line(colour = "black"), 
    panel.border = element_rect(colour = "black", fill = NA, size = 1), legend.position = c(0.8, 
        0.93), legend.title = element_blank())

Fig_Ranges_16S <- grid.arrange(plo_pn_16S, plo_pn_den_16S, plo_habn_16S, top = "ASV range size 16S")
```

```
ggsave("ASV_ranges_16S.pdf", plot = Fig_Ranges_16S, width = 12, height = 8)
```

```
######################################### ASV 'ranges'

head(out_abus_18S)
```

```
## Warning: `...` is not empty.
## 
## We detected these problematic arguments:
## * `needs_dots`
## 
## These dots only exist to allow future extensions and should be empty.
## Did you misspecify an argument?
```

```
## # A tibble: 6 x 7
##   ASV_ID  taxonomy                      site  presence tax.class  habitat sample
##   <chr>   <chr>                         <chr>    <dbl> <chr>      <chr>   <chr> 
## 1 ASV1094 Eukaryota;Amorphea;Obazoa;Op~ TF12S        5 Fungi      TF      S     
## 2 ASV1250 Eukaryota;SAR;Alveolata;Cili~ TF12S        1 Ciliophora TF      S     
## 3 ASV1599 Eukaryota;SAR;Alveolata;Cili~ TF12S        1 Ciliophora TF      S     
## 4 ASV580  Eukaryota;Archaeplastida;Chl~ TF12S        1 Chloropla~ TF      S     
## 5 ASV50   Eukaryota;Amorphea;Obazoa;Op~ TF12S       16 Fungi      TF      S     
## 6 ASV850  Eukaryota;Amorphea;Obazoa;Op~ TF12S        1 Fungi      TF      S
```

```
pn_18S <- out_abus_18S %>% select(-taxonomy, -habitat) %>% group_by(tax.class, ASV_ID) %>% 
    summarise(plot_number = n()) %>% ungroup()
```

```
## `summarise()` regrouping output by 'tax.class' (override with `.groups` argument)
```

```
# number of habitats
hab_n_18S <- out_abus_18S %>% filter(!is.na(habitat)) %>% select(-taxonomy) %>% group_by(tax.class, 
    ASV_ID, habitat) %>% summarise(ASVs.number = n()) %>% ungroup() %>% select(-ASVs.number) %>% 
    group_by(tax.class, ASV_ID) %>% summarise(habitat_number = n()) %>% group_by(tax.class, 
    habitat_number) %>% summarise(num_hab_num = n()) %>% mutate(fraction_hab_num = round(num_hab_num/sum(num_hab_num), 
    3)) %>% ungroup()
```

```
## `summarise()` regrouping output by 'tax.class', 'ASV_ID' (override with `.groups` argument)
## `summarise()` regrouping output by 'tax.class' (override with `.groups` argument)
## `summarise()` regrouping output by 'tax.class' (override with `.groups` argument)
```

```
cols_tax <- c("#8dd3c7", "#ffffb3", "#bebada", "#fb8072", "#80b1d3", "#fdb462", "#b3de69", 
    "#fccde5", "#d9d9d9", "#bc80bd", "#ccebc5")

plo_pn_18S <- ggplot(data = pn_18S, aes(x = tax.class, y = plot_number, fill = tax.class)) + 
    geom_bar(stat = "identity", position = "dodge") + scale_fill_manual(values = cols_tax, 
    name = "") + ylab("Number of plots") + ggtitle("A) Number of plots") + theme_bw() + 
    theme(legend.position = "none") + theme(panel.grid.major = element_blank(), panel.grid.minor = element_blank(), 
    panel.background = element_blank(), axis.line = element_line(colour = "black"), 
    panel.border = element_rect(colour = "black", fill = NA, size = 1))

plo_pn_den_18S <- ggplot() + geom_density(data = pn_18S, aes(plot_number, group = tax.class, 
    col = tax.class), size = 1) + scale_colour_manual(values = cols_tax, name = "") + 
    ggtitle("B) Number of plots - density") + xlab("Number of plots") + ylab("Density") + 
    xlim(c(1, 39)) + guides(color = guide_legend(ncol = 2)) + theme_bw() + theme(panel.grid.major = element_blank(), 
    panel.grid.minor = element_blank(), panel.background = element_blank(), axis.line = element_line(colour = "black"), 
    panel.border = element_rect(colour = "black", fill = NA, size = 1), legend.position = c(0.65, 
        0.78), legend.title = element_blank())


plo_habn_18S <- ggplot(data = hab_n_18S, aes(x = tax.class, y = fraction_hab_num, 
    fill = as.factor(habitat_number))) + geom_bar(stat = "identity", position = position_dodge()) + 
    scale_fill_brewer(palette = "Blues", name = "") + ylab("Fraction of ASVs") + 
    ggtitle("C) Number of Flood Class") + guides(fill = guide_legend(ncol = 4)) + 
    theme_bw() + theme(panel.grid.major = element_blank(), panel.grid.minor = element_blank(), 
    panel.background = element_blank(), axis.line = element_line(colour = "black"), 
    panel.border = element_rect(colour = "black", fill = NA, size = 1), legend.position = c(0.8, 
        0.93), legend.title = element_blank())

Fig_Ranges_18S <- grid.arrange(plo_pn_18S, plo_pn_den_18S, plo_habn_18S, top = "ASV range size 18S")
```

```
ggsave("ASV_ranges_18S.pdf", plot = Fig_Ranges_18S, width = 12, height = 8)
```

```
## Summarizes data.  Gives count, mean, standard deviation, standard error of the
## mean, and confidence interval (default 95%).  data: a data frame.  measurevar:
## the name of a column that contains the variable to be summariezed groupvars: a
## vector containing names of columns that contain grouping variables na.rm: a
## boolean that indicates whether to ignore NA's conf.interval: the percent range
## of the confidence interval (default is 95%)
summarySE <- function(data = NULL, measurevar, groupvars = NULL, na.rm = FALSE, conf.interval = 0.95, 
    .drop = TRUE) {
    library(plyr)
    
    # New version of length which can handle NA's: if na.rm==T, don't count them
    length2 <- function(x, na.rm = FALSE) {
        if (na.rm) 
            sum(!is.na(x)) else length(x)
    }
    
    # This does the summary. For each group's data frame, return a vector with N,
    # mean, and sd
    datac <- ddply(data, groupvars, .drop = .drop, .fun = function(xx, col) {
        c(N = length2(xx[[col]], na.rm = na.rm), mean = mean(xx[[col]], na.rm = na.rm), 
            sd = sd(xx[[col]], na.rm = na.rm))
    }, measurevar)
    
    # Rename the 'mean' column
    datac <- rename(datac, c(mean = measurevar))
    
    datac$se <- datac$sd/sqrt(datac$N)  # Calculate standard error of the mean
    
    # Confidence interval multiplier for standard error Calculate t-statistic for
    # confidence interval: e.g., if conf.interval is .95, use .975 (above/below), and
    # use df=N-1
    ciMult <- qt(conf.interval/2 + 0.5, datac$N - 1)
    datac$ci <- datac$se * ciMult
    
    return(datac)
}
```

```
###################################################################### Plots taxonomic groups############################

head(FunGuild_abus_18S)
```

```
##    ASV_ID          Guild  site presence    guild.class habitat sample
## 1 ASV1094       Parasite TF12S        5       Parasite      TF      S
## 2   ASV50     Saprotroph TF12S       16     Saprotroph      TF      S
## 3  ASV850        unknown TF12S        1        unknown      TF      S
## 4  ASV240        unknown TF12S        2        unknown      TF      S
## 5   ASV99       Parasite TF12S        1       Parasite      TF      S
## 6 ASV1321 plant_pathogen TF12S        1 plant_pathogen      TF      S
```

```
# load 18S
hab_guild_18S <- FunGuild_abus_18S %>% distinct(ASV_ID, site, habitat, .keep_all = T) %>% 
    group_by(site, habitat, guild.class) %>% dplyr::summarise(ASVs.number = n()) %>% 
    mutate(fraction.ASVs = round(ASVs.number/sum(ASVs.number), 3))
```

```
## `summarise()` regrouping output by 'site', 'habitat' (override with `.groups` argument)
```

```
ord <- c("TF", "HV", "MV", "LV")
hab_guild_18S$value <- factor(hab_guild_18S$habitat, levels = ord)

# number

cols <- c("#238b45", "#c7e9c0", "#74c476", "#00441b")

hab_guild_18S2 <- summarySE(hab_guild_18S, measurevar = "ASVs.number", groupvars = c("guild.class", 
    "habitat"))
```

```
## ------------------------------------------------------------------------------
```

```
## You have loaded plyr after dplyr - this is likely to cause problems.
## If you need functions from both plyr and dplyr, please load plyr first, then dplyr:
## library(plyr); library(dplyr)
```

```
## ------------------------------------------------------------------------------
```

```
## 
## Attaching package: 'plyr'
```

```
## The following object is masked from 'package:ggpubr':
## 
##     mutate
```

```
## The following object is masked from 'package:matrixStats':
## 
##     count
```

```
## The following object is masked from 'package:IRanges':
## 
##     desc
```

```
## The following object is masked from 'package:S4Vectors':
## 
##     rename
```

```
## The following objects are masked from 'package:dplyr':
## 
##     arrange, count, desc, failwith, id, mutate, rename, summarise,
##     summarize
```

```
## The following object is masked from 'package:purrr':
## 
##     compact
```

```
## Warning in qt(conf.interval/2 + 0.5, datac$N - 1): NaNs produced
```

```
# The errorbars overlapped, so use position_dodge to move them horizontally
pd <- position_dodge(0.1)  # move them .05 to the left and right

Hab_N_18S_Guild <- ggplot(data = hab_guild_18S2, aes(x = guild.class, y = ASVs.number, 
    fill = habitat)) + geom_bar(stat = "identity", position = "dodge") + scale_fill_manual(values = cols, 
    name = "") + geom_errorbar(aes(ymin = ASVs.number - se, ymax = ASVs.number + 
    se), position = position_dodge()) + scale_color_manual(values = cols, name = "") + 
    ggtitle("Funal functional guilds") + ylab("Number of ASVs") + xlab("") + coord_flip() + 
    theme(panel.grid.major.x = element_line(size = 0.5, colour = "#d9d9d9"), panel.grid.major.y = element_blank(), 
        panel.grid.minor.x = element_line(size = 0.5, colour = "#f0f0f0"), panel.grid.minor.y = element_blank(), 
        panel.background = element_blank(), axis.line = element_line(colour = "black"), 
        panel.border = element_rect(colour = "black", fill = NA, size = 1), legend.position = "right")


ggsave("figGuildMean3.pdf", plot = Hab_N_18S_Guild, width = 12, height = 8)
Hab_N_18S_Guild
```
