## Appendix5 for "Above- and below-ground biodiversity responses to the prolonged flood pulse in central-western Amazonia, Brazil": paper2_Appendix5.html

Javascript must be enabled to view this page.

magnitude
magnitudeUnassigned

jurua16s---ssu---krona---Tax\_16S\_Silva2----Total---sim\_93---tax\_silva---td\_20

10211

129

1

1

1

19

9

10

9

1

1

10

10

10

10

10

18

18

15

3

2

2

1

81

2

2

1

1

1

50

50

50
1

25

24

29

29

14

14

4

4

11

11

10057

6

6

6

5

1

1

3

3

1

1

1

1

1

1

88

1

1

10

10

65

2

2

6

6

1

2

1

1

1

17

1

3

1

1

1

12

1

4

6

1

1

1

1

1

1

5

1

1

4

1

1

1

1

4

29

5

1

4

23

6

1

1

1

3

1

1

1

2

2

2

1

1

1

1

7

7

7

7

1

1

1

1

4

1

1

3

3

2

1

2

1

1

1

1

1

1

1

1

289

240

240
198

9

3

2

1

8

22

25
7

2

4

2

2

2

6

1
3

2

10

2

2

2

6

1

4

1

1

84

43

43

16

16

1

6

9

27

8

8

10

10

2

7

7

1308

4

1

193

10

10

145

14

10

1

3

125

4

3

118

6

6

38

37

1

91

1

8

34

65

10

16

1

10

1

1

1

1

1

1

4
341

15

7

7

8

142

142
4

1

1

82

2

45

7

46

6

1

24

5
24

2

17

1

102
46

55
56

1

544

1
520

5
423

30

8

15

129

18

45

20

9

120

19

5

96

2

22

49

49

49

49

1

41

7

1767

1

1

2

23

16

1674

4

307

307

307

243

413

413

413

1

15

3

7

7

674
2

428

1
187

16

2

33

14

33

24

5

8

51

57

57

51

37

37
4

8

12

9

4

14

2845

24

382

157
2

41

27

27

12

5

7

75
1

20

8

46

85

140

1
2434

125

1
125

9

102

13

276

276
3

10

6

12

2

14

6

4

118

5

5

31

22

2

2

3

8

23

176

2
23

1

4

2

1

2

1

4

6

9

4

4

1

2

2

112
2

1

4

1

1

14

5

1

3

12

1

4

1

15

2

10

1

4

3

2

25

1

1

1

1

6

2

2

2

18

2

3

3

1

2

4

3

1

1

3

1

2

38

24

24

14

14

255

5

1

1

2

1

250

215

22

5

6

1

1

2

2

2

43

1
36

11

11

1

12

7

3

1

1

2

136

136

1

25

85

12

13

755
6

632

632

79

3

76

1

1

8

8

17

4

5

8

4

4

8

1

239

186

186

47

9

5

33

2

2

4

4

2

385

385
10

9

2

21

1

9

4

3

7

27

47

2

2

1

1

4

5

7

12

2

41

12

120

1

4

29

1

1

3

1

1

1

1

1

2

2

2

12

12

12

12

12

8

105

4

11

11

1

67

2
67

58
65

7

22

22
10

12

12

1

3

2

2

1

875

253

1

63

63

2

1

7

48

2

3

2

1

1

186

164

13
22

9

1

621

8

433

433
3

2

1

17

312

98

12

12

1

6

5

8

8

6

1

1

160

160

93

3

2

26

34

2

2427

8

1

1

1

7

7

7

2419

156

156
4

7

2

3

4

17

1

1

4

7

1

3

96

6

1

1

17

1

5

10

1

4

2

2

2

1

1

2

2

1

1

2

1

1

3

3

1

2

48

213

6

194

13

1

11

1

45

45

7

38

331

1
233

5

113

26

16

50

22

98

23

62

13

41

20

1

1

11

11

9

3

6

383

4
383

44

1

65

2

1

31

13

1

191

10

2

2

4

12

17

11

11

1

1

5

1

2

1

1

8

8

1

1

1

2

1

2

140

4
140

4

10

122

39

39

39

2

2

2

7

6

1

1

949

15

15

184

10

3

27

9

1

1

4

58

9

12

22

2

7

16

3

1

1

18

12

49

24

1

2

22

23

11

2

1

1

2

138

4

1

5

2

65

6

1

9

2

5

7

1

1

3

26

3

12

11

1

1

1

8
363

15

2

1

9

2

1

7

6

25

1

1

67

215

3

34

34

43

8

35

38

3

5

19

10

1

10

1

1

9

73

1

24

8
4

4
3

1

16

16

48

30

1

7

7

1

1

1

1

2

1

10

1

1

1

1

3

1

1

1

1

4

4

16

12

12

12

12

4

4

42

1

1

41

41

41

37

4

6

6

6

6

25
