## Appendix6 for "Above- and below-ground biodiversity responses to the prolonged flood pulse in central-western Amazonia, Brazil": paper2_Appendix6.html

Javascript must be enabled to view this page.

magnitude
magnitudeUnassigned

jurua18s---ssu---krona---Jurua\_18S----Total---sim\_90---tax\_silva---td\_20

2052

157

1895
1

93

93

5

1
2

1

1

3

3

3

88

1

1

1

87

1

1

1

86

86

80

1

1

1

1

1

1

1

78

6
77

8

5

1

2

1

1

1

1

1

1

1

1

2

2

9

3

1

1

1

1

1

1

1

1

1

2

1

1

1

1

8

1

1

1

1

1

1

1

1

11

2

1

2

1

1

2

1

1

1

1

1

1

1

1

4

1

1

1

1

5

1

1

2

1

1

1

2

1

1

2

2

1

1

1

1

6

1

1

1

2

1

4

4

1

1

1

1

3

1

1

2

2

2

1

997

904

904

36

36

10

2

1

1

1

1

1

8

8

8

2

2

2

2

2

1

1

1

1

1

6

2

2

2

2

2

4

3

3

3

3

3
2

1

1

1

1

1

1

1

1
26

2

1

1

1

1

1

17

1

3
15

1

1

4

3

1

7
2

2

3

1

1

1

6

6

6
3

3

868

8

8

8

1
3

1

1

857
2

61

28

28

12

1

1

1

1

1

1

2

1

1

4

1

2

1

1

1

1

1

1

13

13

9

4

3

3

3

11

11

1
11

8

8

2

22

22
2

3

2

2

1

1

14

13

6

7

1

3

1

2

2

6

1

1

1

1

1

1

621

375

19

19

19

4

1

2

1

4

1

1

1

1

4

1

1

2

4

1

2

1

1

1

1

1

1

1

1

1

1

1

1

355
4

2
19

5

4

4

1

4

4

4

3

1

1

2

2

4

2
4

1

1

1

1

1

4

4

1

1

3

3

3

3

3

3

1

1

1

1

7
132

17
9

1

1

3
7

2

2

10

5

5

5

5

10
3

7

1

1

2

2

1

1

1

1

7

2

1

1

4

1

2

1

22

1
3

1

1

19

19

4

4

4

1

1

1

2

1
2

1

1

1

1

3

3

1

2

6

6

4

2

38

1

2

4

3

1

3
2

1

6

6

1

7

7

1

5

4

1

2

1

1

6

2

2

2

3

2

1

1

1

1

7

3

3

3

1

1

1

5

3

2

1

1

1

2

10

10

1

1

1

3

1

2

4

1

1

2

1

1

78

3
42

9

2

5

2

4

4

1

1

6

19

15

1

3

2

2

2

6

5

2

3

1

1

1

1

1

21

7

6

1

14

7

7

1

1

1

3

1

1

2

2

2

2

1

1

1
76

1

1

1

1

9

9

1

8

7

1

1

1

1

2

2

2

18

1

1

1
2

1

4

1

4

4

2

2

2

1

1

2

17
5

2
12

2

5

3

14

1
10

2

2

5

4

4

5

5

2

2

2

1

1

1

5
12

2
6

2

2

2

1

1

1

1

1

246

215

1

1

1

1

15

3

3

1

1

1

1
12

1

1

3

2

1

1

1

1

1

2

2

1

1

1

1

199
15

1

1

1

23

7
23

3

5

2

1

2

1

2

2

2

2

3

3

2

1

4

1

1

2

2

1

10
1

1

1

1

1

2

2

1

1

1

3

1

2

1
8

2

2

4

4

1

1

14

1

1

1

3

2

1

2

4

1

2

1

3

6
94

2
16

2

1

1

2

1

4

3

1

1

2

2

1

6

6

1

1

1

1

1

2

2

7

1

4

2

3

2

1

1

8

8

5
11

2

2

2

1

1

6

5

1

1

2

4

1

1

2

10

2

1

1

8

4

4

2

2

2

1

1

1

3

3

3

13

23
2

6

5
1

2
4

1

1

1

1

1

4

4

4

1

2

1

2

2

1

1

6

4

4
1

1

1

1

2

2

2

2

2

2

1

8

3

3

3

3

5

1
4

3
2

1

1

1

2
38

7

6

6

1

1

5

5

1

1

1

1

7

7

1

1

1

6

6

3

1

1

1

22

22

22
15

1

1

2

2

2

38

13

13

13

13

4

9

22

3

12

12

12

12

3

4

3

1

5

1

4

11

67

67

4
56

5

2
5

3

1
18

5
2

3

5

2

4
5

1

2

1

1

5

4

4

1

1

6

2

2
3

1

1

1

1

1

1

13

6

2

3

1

4
5

1

2
1

1

1

1

1

11

11

1

1

7

7

3

3

93
13

6

8
1

2

2

1

2

1

1

8

2

1

1

1

3
1

2

2

3

3

6

4

4

1

3

2

2

2

1
38

3
14

2

2

9

3
12

1

2

6

11
8

1

2

8
1

3

4

1

1

1

1

4

3

3

1

1

1

1

797

90

48

5

5

1

1

1
4

1

2

40

1

1

12

4

4

3

1

1

1

4

20

5

1

1

9

2

2

1

1

1

1

2

2
1

1

5

5
3

2

3

3

13

4

5

3

1

1

3
16

2

1

2

1

1

1

2

3

4

1

1

1

1

2

381

277

4

4

3

1

273

3
76

20
67

2

1

1

1

1

2

4

1

2

1

2

3

8

1

1

1

1

5

5

1

2

1

4

2

2

139

4

3

1

64

13
36

2

1

3

1

12

2

2

1
3

2

5

5

14
1

1

11

1

6

2

2

2

3
12

2

3

4

56

3

3

6

4

2

38
7

4

3

4

3

5

1

5

1

2

3

6
1

1

2

1

1

2

1

1

1

1

11

10
7

3

1

3
47

8
43

2

2

1

2

7

1

1

1

1

4

3

3

5

2

1

1

2

2

12

10

6

1

1

2

1

1

1

1

2

1

1

3

2

1

1

1

1

1

1

1

1

22

3

3

5

5

12

3

1

4

1

2

1

2

2

1

67

67

9

2
6

2

1

1

3

1

2

58

47
2

6

1

1

1

17

6

1

1

1

1

7

2

2

1

1

3

6

6

326

1

1

1

1

6
325

2

1

1

65
13

2

1

21

3

25

1

1

1

1

1

1

1

1

44
8

8

3

1

2

2

2

3

3

25

3

3

18

9

9

5

69

59

48
2

3

1

2

16

16

9

18

15

1

2

11

6
3

1

2

5
1

1

1

1

1

5

5

4

1

1

1

30
1

9

2

6

12

6

1

3

3

2

2

36

1

13
32

1

18
